# Global changes in unproductive splicing and the NMD system efficiency in tumors

**DOI:** 10.64898/2025.12.10.692474

**Authors:** L.G. Zavileyskiy, A.A. Mironov, D.D. Pervouchine

## Abstract

The Nonsense-Mediated Decay (NMD) pathway is a mRNA quality control mechanism, which not only degrades deleterious transcripts but also orchestrates a large number of post-transcriptional regulatory programs through unproductive splicing. Here, we developed a robust metric derived from the quantification of splicing in the RNA-seq data to measure NMD efficiency at a sample level. We demonstrate that NMD efficiency varies substantially both between and within tissues, with the magnitude of variation comparable to that observed upon knockdown of the core NMD factor UPF1. By analyzing TCGA cancer cohorts, we further show that, in many tumors, unproductive splicing events exhibit coordinated changes towards either collective suppression or collective activation of NMD isoforms, which is indicative of a global deregulation of the NMD pathway activity. Consistently, we observed a striking divergence of NMD efficiency in cancers from the tissue-specific baseline level suggesting that tumors partially erase the NMD signature of their tissue of origin. The application of the developed metric to RNA-binding protein knockdowns allowed identification of several novel potential regulators of NMD efficiency. In sum, this study provides a solid framework for NMD efficiency quantification, describes its biological and clinical relevance, and opens new avenues for dissecting mechanisms of post-transcriptional gene expression regulation by the NMD pathway.

## Introduction

The nonsense-mediated mRNA decay (NMD) pathway is a mRNA surveillance system that selectively eliminates transcripts containing premature termination codons [1]. Such transcripts may arise not only as a result of nonsense or frameshift mutations but also as a result of alternative splicing [2]. On the one hand, degradation of mRNAs with premature termination codons prevents the production of deleterious proteins. On the other hand, beyond this classical role, NMD also acts together with alternative splicing as a widespread post-transcriptional mechanism of gene expression regulation, referred to as unproductive splicing [2, 3]. The activity of the NMD system exerts a substantial impact on the transcriptome composition and has been linked to diverse physiological processes and pathological conditions such as cellular differentiation, lymphocyte development, stress response, genetic disorders, and cancers [4, 5, 6, 7, 8, 9].

Despite the existing knowledge, the role of NMD in tumorigenesis remains controversial [10, 8, 9]. While cancer cells may exploit NMD to downregulate tumor suppressors, their adaptation to stress conditions such as hypoxia or therapeutic pressure often requires precise tuning of its efficiency [11, 12]. As a result, NMD can exert both pro- and anti-tumorigenic effects depending on the conditions and the microenvironment [13, 14, 15, 12, 16]. In accordance with this, the core NMD factor UPF1 has been reported to be downregulated in certain cancers and upregulated in the others [17, 18, 19, 20]. Measuring sample-level NMD efficiency is challenging partly because NMD factors themselves undergo negative autoregulation, making their mRNA levels unreliable proxies for the pathway activity [21, 22].

Transcript-level determinants of NMD efficiency have been widely studied [11, 23, 24], yet the information on global, condition-specific NMD efficiency often remains contradictory. Early works used reporter constructs and monitored a small number of endogenous NMD targets, revealing apparent inter-tissue variability, but these estimates could suffer from gene-specific regulatory biases [25, 22]. Approaches based on allele-specific expression of transcripts carrying nonsense mutations concluded that NMD efficiency is largely uniform across tissues [23, 24], but a later study of expression levels of NMD-sensitive isoforms and matched productive isoforms again reported non-random inter-tissue differences [26].

Motivated by this conflicting evidence, we sought to develop a robust and sensitive metric for quantitative assessment of NMD efficiency based on RNA sequencing (RNA-seq) data. Here, we introduce an integral NMD efficiency metric based on unproductive splicing events (USE), i.e. local alternative splicing events that generate isoforms containing premature termination codons and are sensitive to NMD inhibition. USEs are far more abundant than heterozygous nonsense mutations and provide substantially greater statistical power for sample-level estimates. Estimates using this metric confirmed that inter-tissue differences may reach up to 80% of the effect size observed in UPF1 knockdown. The application to cancer data demonstrated a correlation between the direction of change in NMD efficiency in tumors and the direction of change in most individual events. The metric further showed that tumors frequently lose the NMD efficiency levels characteristic of their tissue of origin — an alteration that, in several cancer types, correlates with unfavorable prognosis. Juxtaposition of USE splicing changes with corresponding gene-expression changes allowed to distinguish event-specific splicing regulation from global NMD efficiency shifts. By applying the developed metric to RNA-binding protein (RBP) knockdown datasets, we recovered known NMD factors and proposed new candidate regulators of the NMD pathway.

## Materials and Methods

### RNA sequencing data

Transcriptomes of healthy and tumor human tissues from The Cancer Genome Atlas (TCGA) were downloaded from the dbGaP portal in the form of alignments to the GRCh38 human genome assembly. Thirteen tumor cohorts containing at least 15 paired tumor-normal samples were selected (Table S1). In differential splicing analysis, only matched tumor-normal samples were used. Transcriptomes of human tissues from the Genotype Tissue Expression (GTEx) V7 project were downloaded in FASTQ format from the dbGaP portal. We selected 10,100 samples with a read length of 75 nt, containing at least 20 million reads per sample (Table S2). Transcriptomes of K562 and HepG2 cells subjected to RBP knockdowns or knockouts were downloaded from the ENCODE consortium website [27] in the form alignments to the GRCh38 human genome assembly. Perturbations performed in two biological replicates in one or both cell lines were selected (Table S3). Transcriptomes of HeLa cells subjected to knockdowns of spliceosomal components were obtained from Array Express under accession number E-MTAB-11202 and converted to FASTQ format [28]. Each knockdown was represented by one biological replicate. Transcriptomes of HeLa cells subjected to knockdowns of UPF1, SMG6, SMG7, and double knockdown of SMG6 and SMG7, as well as rescue experiments, were obtained from the SRA repository under the accession number GSE86148 [29]. Reads from the samples in FASTQ format were mapped to the human genome version GRCh38 using STAR aligner v2.7.8a [30] using GENCODE 43 [31] annotation as a reference. Gene expression levels were assessed using the FeatureCounts utility [32]. Differential gene expression in TCGA was analyzed using limma-voom from the edgeR package [33]. In RBP and NMD system inactivation experiments, differential expression was analyzed using DESeq2 [34].

### USE Catalog

A catalog of annotated unproductive splicing events (USEs) was generated using NMDj utility [35] based on the ENSEMBL genome annotation (version 108). To identify novel USEs, transcriptomes of each TCGA sample were assembled using StringTie v2.2.1 software with the –conservative option [36], and the aggregated annotation across all samples was fed to NMDj. Additionally, skipping events of all constitutive exons flanked by constitutive introns were added to the list of novel events. Novel USEs whose characteristic intron sets of NMD transcripts overlapped with the intron sets of annotated USEs were removed in order to obtain non-redundant catalogs.

### Quantitative assessment of splicing

The number of split reads supporting intron splicing was computed from short read alignments using the IPSA package with the default settings [37]. The splicing rate of each USE was characterized by the T metric calculated using the NMDj package [35], defined as the ratio of the number *a* of split reads supporting the NMD isoform to the total number *a* + *b* of split reads supporting both NMD and the coding isoforms (the value T = 0 indicates the absence of the NMD isoform). Only USEs with *a* + *b* > 15 in at least half of the samples in each comparison group were considered. Differential splicing analysis in TCGA and NMD inactivation experiments was estimated using a random-effects model implemented in the statsmodels.stats.meta analysis module. The arcsine transformation was applied to T values to stabilize variance. The within-sample variance *s* was estimated using the formula 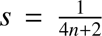, where *n* = *a* + *b* [38], and the between-sample variance was estimated using the non-iterative method of DerSimonian-Laird [39]. Estimates of the group means and total variances were used to conduct *z*-tests for the significance of differences between groups. For RBP perturbation experiments with few or no biological replicates, *a* and *b* values were pooled across all replicates, and the statistical significance of T differences between groups was assessed using the proportion test. To compare splicing rates between knockdown and control conditions, ΔT = T*_KD_* − T*_C_* was used, where T*_KD_* and T*_C_*are the T values in knockdown and control, respectively. ΔT values were averaged across all inactivation experiments of the same RBP, and *P*-values were calculated using the Stauffer *z*-score method [40].

### NMD Efficiency Assessment

To select baseline USEs that specifically respond to the inactivation of the NMD system, but not to that of individual RSBs, the following metric α was used. This metric was calculated as the product of the ΔT value between the RBP inactivation and the control, normalized by the standard deviation across all RBPs, and the *z*-score of the corresponding *P*-value. It was required that the ratio of the α value for NMD inactivation experiments to the largest α value for all RBP from the panel of perturbation experiments was not less than *L* (see below). For each data set (TCGA, GTEx, ENCODE), core USEs were selected from the set of events, for which the proportion of samples with *a* + *b* > 15 was at least *N* (see below), and the proportion of samples with 0 < T < 1 was at least *K* (see below). For each core event, the T values were transformed to *z*-scores across all samples. The NMD efficiency metric of a sample was defined as the negative mean of *z*-transformed T values across all core events.

The parameters *N*, *K*, and *L*, as well as core USEs, were optimized separately for each dataset due to biological (tumor, tissue, perturbation) and technical (sequencing depth) heterogeneity. For TCGA: *N* > 0.5, *K* > 0.1 in at least 12 of 13 cohorts, *L* > 0.3, 120 events were selected. For GTEx: *N* > 0.9, *K* > 0.9 across all tissues, 111 events were selected. For ENCODE: *N* > 0.7, *K* > 0.7 across all experiments, *L* > 0.2, 127 events were selected. Experiments in which less than 70% of baseline USEs met these criteria were excluded from further analysis.

### Survival analysis

Patient survival data were obtained from the UCSC Xena platform [41]. The association between NMD efficacy and overall survival was assessed using the Cox proportional hazards model [cox]. Within each tumor cohort, patients were stratified into two equal groups, with high and low NMD efficacy. Gender and age were included as covariates; age was divided into three equal-sized groups within each cohort.

### NMD efficiency and regulated nonproductive splicing

To assess the abundance of specifically regulated USEs, the following odds ratios were calculated. For a given threshold value of the absolute value of ΔT in tumors, the ratio of the number of USEs demonstrating opposite directions of splicing and expression changes and the number of USEs with co-directional changes among events with |ΔT| above and below the threshold value were compared. Only genes with | log_2_ *FC*| > 0.3 and *P* < 0.05 were considered. The same estimates were calculated separately for positive and negative ΔT values to assess the dependence on the direction of NMD efficiency change.

### Statistical Analysis

All statistical tests were performed in Python 3.9.13. Bonferroni-Holm corrections for multiple testing were performed for differential expression and splicing analyses in TCGA and for RBP or NMD factor inactivation experiments separately for each tumor cohort or experiment, when assessing the significance of differences in NMD efficiency between tumor and normal tissue, when searching for associations of NMD efficiency with patient survival, and when assessing the enrichment of functional categories of RBPs using the GSEA method in a list of RBPs ranked by the magnitude of change in NMD efficiency in the corresponding knockdown. Survival analysis was performed using the lifelines module. Boxplot whiskers in all figures correspond to the extreme points deviating from the median by no more than 1.5 interquartile range units. Error bars in figures 1B, 4, and S3 correspond to 95% confidence intervals. In this paper, *r* and *P* denote the Pearson correlation coefficient and adjusted *P*-value, respectively; FC denotes the fold change in gene expression level. All nonparametric tests were performed using a normal distribution with continuity correction.

**Figure 1:**
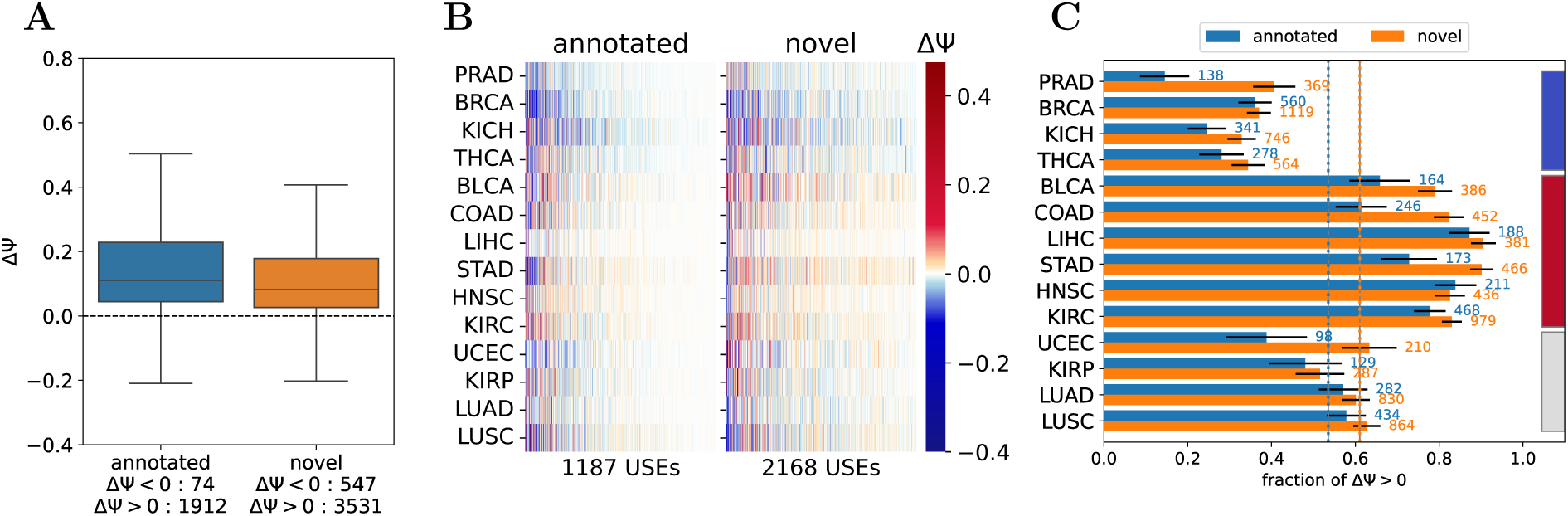
Changes in unproductive splicing upon NMD inactivation and in cancer cohorts from TCGA. **(A)** Changes in splicing rates (ΔT) of USE in double knockdown of SMG6 and SMG7 in the HeLa cell line. **(B)** Changes in splicing rates (ΔT) of USEs that are differentially spliced in at least one TCGA cohort. **(C)** The proportions of USEs with elevated NMD isoforms in tumor cohorts (ΔT > 0). Numbers to the right of the bars indicate the number of USEs. The dotted lines show the average proportions of USE with elevated NMD isoforms across all cohorts. Error bars represent 95% confidence intervals. Blue, red, and gray colors represent tumor groups with the same direction of splicing alterations.

## Results

### Unproductive splicing changes exhibit a preferred cancer-specific direction

Our initial goal was to characterize global unproductive splicing changes in tumors. Using the NMDj tool [35], we assembled a catalog of unproductive splicing events (USE) that were either present in the ENSEMBL genome annotation (annotated) or inferred from RNA-seq data for tumor transcriptomes (novel). We identified 1986 annotated and 4078 novel USEs that respond significantly (*P* < 0.05) to NMD inhibition in the SMG6-SMG7 double knockdown experiments [29]. As expected, for the vast majority of events (more than 75%) the relative abundance of the NMD isoform, as measured by the change in the percent-spliced-in (ΔT, NMD inhibition vs. control) metric, increased (Figure 1A).

USEs that changed in the expected direction, i.e., increased the proportion of NMD isoform under NMD inactivation, were selected for further analysis. We found 1187 annotated and 2168 novel events that were differentially spliced in at least one tumor cohort (*P* < 0.05) with remarkably diverse patterns of both positive and negative changes (Figure 1B). Next, we quantified the proportion of annotated and novel USEs with positive ΔT and tested statistically whether they differ from the respective average ΔT value across all cancer cohorts (Figure 1C). Interestingly, most cancer types exhibited a preferred direction of unproductive splicing change: in PRAD, BRCA, KICH, and THCA, the proportion of NMD isoforms tends to decrease, whereas in BLCA, COAD, LIHC, STAD, HNSC, and KIRC it more often increases. This trend was observed for both annotated and novel events; however, novel events showed a higher fraction of positive changes, consistent with their absence from genome annotations due to low expression levels in normal conditions.

### USE-based metric robustly estimates NMD efficiency

The consistency in the direction of unproductive splicing changes across tumors suggests that these alterations may be driven by global changes of the NMD pathway activity. To estimate NMD efficiency, we selected a core set of *n* = 120 USEs that (i) were expressed in at least 13 of the 14 analyzed TCGA cohorts, (ii) responded to NMD inhibition in the expected direction, i.e., by upregulation of NMD-sensitive transcripts, and (iii) were not sensitive to perturbations of RBP expression levels (in order to reduce the confounding effect of event-specific regulation, see Methods). For each sample, the NMD efficiency score was defined as the negative mean of *z*-transformed T values of the selected core events.

We validated this metric in several ways. First, the NMD efficiency scores were robust with respect to choosing a core set of events provided that it was large enough (*n* > 100) (Figure S1, see Methods). Second, all USE in the core set showed highly coordinated splicing changes across TCGA samples (Figure S2). Third, the NMD efficiency score dropped in single and double knockdown experiments of three key NMD factors, UPF1, SMG6, and SMG7, and recovered in the respective rescue experiments (Figure 2A). Next, changes of NMD efficiency score in tumors vs. matched normal tissues positively correlated with changes in UPF1 expression levels (Figure 2B). As expected, the fraction of USE with collective positive ΔT changes in TCGA samples dropped from 80% to 20% with increasing the NMD efficiency score (Figure 2C). Finally, applying our metric to healthy tissues from the GTEx project revealed a strong correlation with an existing allele-specific estimate of NMD efficiency [23], in which the latter explained 62% of the variance of the former if brain samples were excluded (Figure 2D). Notably, brain tissues deviated from the general linear trend in such a way that allele-specific estimates for different brain subregions were highly variable, while NMD efficiency score introduced here had a consistent large value.

**Figure 2:**
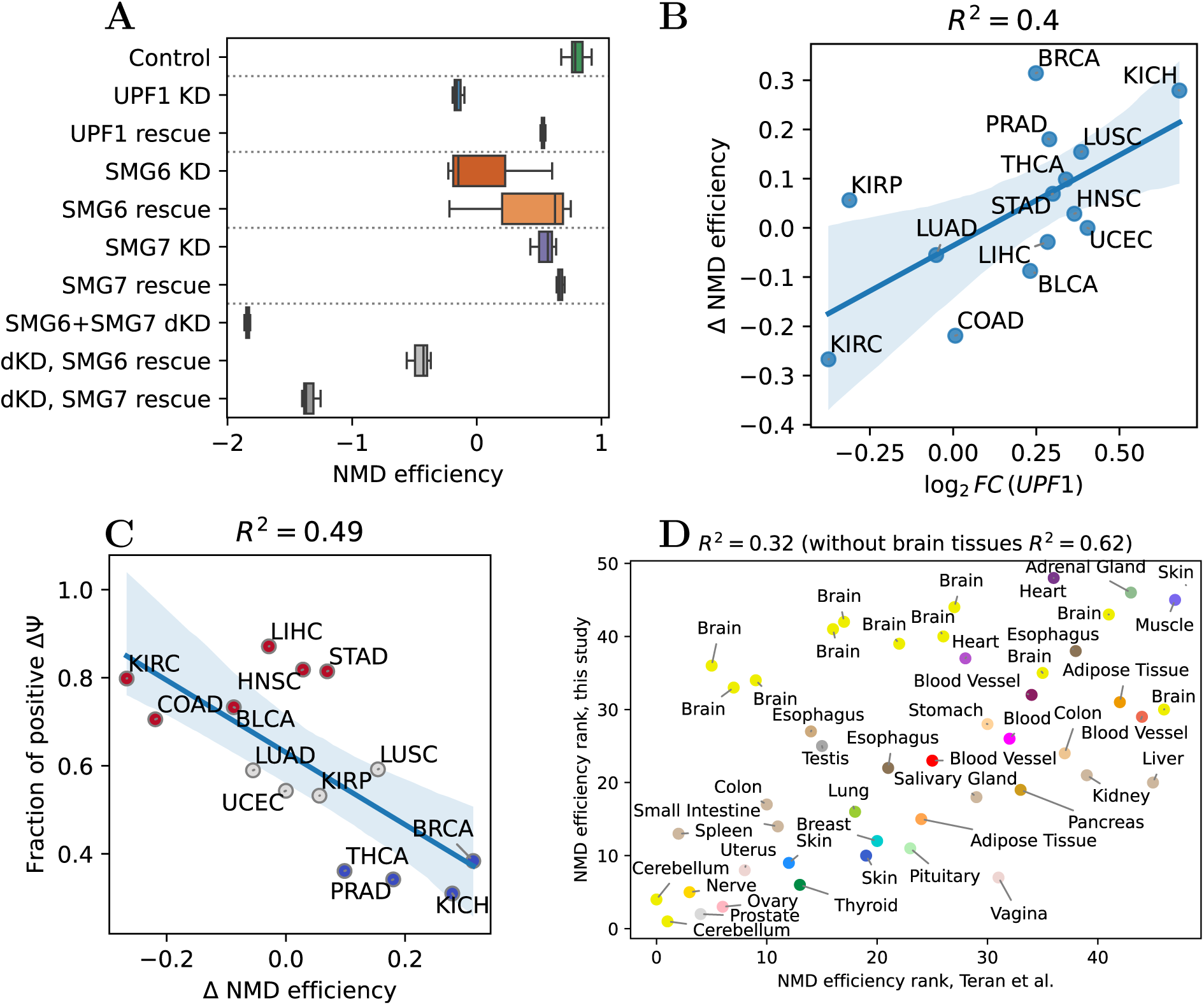
The NMD efficiency estimate. **(A)** NMD efficiency upon knockdown and rescue of NMD factors expression in the HeLa cell line. **(B)** Correlation of changes in NMD efficiency and changes in UPF1 expression levels in tumors. **(C)** Correlation of changes in NMD efficiency and proportion of USEs with positive splicing changes in tumors. Dot colors correspond to the colors of the tumor groups with the same direction of splicing changes in Fig. 1C. **(D)** Correlation of the NMD efficiency score in GTEx tissues from this study with the score based on nonsense mutations [23].

### NMD efficiency varies substantially both between and within human tissues

While the existing NMD efficiency metrics allow relative comparisons across samples, they do not provide a sense of its absolute magnitude [23, 24, 26]. To address this, we introduced a reference scale by comparing the observed NMD efficiency variation to the effect size for the knockdown of UPF1, the core factor of NMD, for which the expression level dropped 3.5 times [29]. With this scaling, the NMD efficiency score in unperturbed conditions was assumed to be 0, while under UPF1 depletion it was assumed to be −1.

In application to the GTEx dataset, this transformed metric confirmed that NMD efficiency varies in a non-random manner across tissues, consistently with recent findings [26] (Figure 3). Of note, 46% of the variance was explained by the tissue factor (one-way ANOVA, *P* ≈ 0), and the difference between tissues with the highest (heart) and the lowest (cerebellum) median NMD efficiency score was comparable to 80% of the effect size of UPF1 knockdown. The variability within tissues was also remarkably high, with interquartile ranges reaching up to 40% of the effect size of UPF1 knockdown in heart, uterus, liver and kidney. The least variable NMD efficiency levels were observed in brain subregions, but the difference between median NMD efficiency levels in the brain cortex and in the cerebellum was about a half of the effect size of the UPF1 knockdown, confirming that these two areas are drastically different in terms of NMD activity [26].

**Figure 3:**
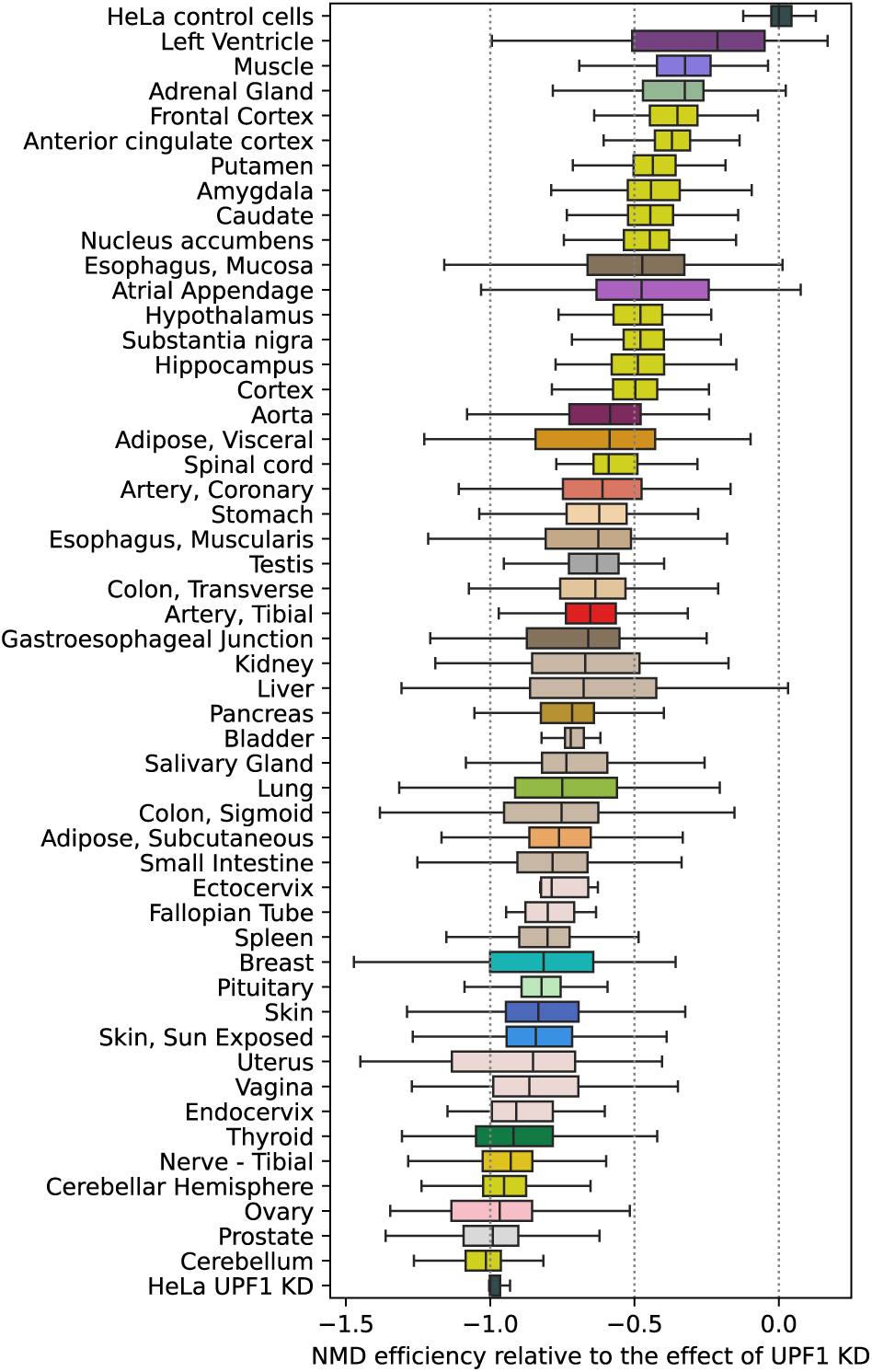
The distribution of NMD efficiencies in GTEx tissues relative to the magnitude of the effect of UPF1 knockdown. NMD efficiencies in control and UPF1 knockdown HeLa cells were set to 0 and −1, respectively.

### NMD efficiency is associated with patient survival and regulated unproductive splicing

In order to examine the variability of NMD efficacy across tumors, we analyzed TCGA cohorts and found significant changes between NMD activity in cancer and the respective normal tissue in BRCA, PRAD, KICH, and KIRC (Figure 4A). Remarkably, the direction of NMD efficiency change in cancer depended on the baseline efficiency of the corresponding normal tissue: in tissues with low baseline NMD efficiency it tends to increase, whereas in tissues with high baseline NMD efficiency it tends to decrease. This led us to a non-trivial conclusion that tumors show a consistent trend toward erasing the tissue-specific NMD efficiency signature.

**Figure 4:**
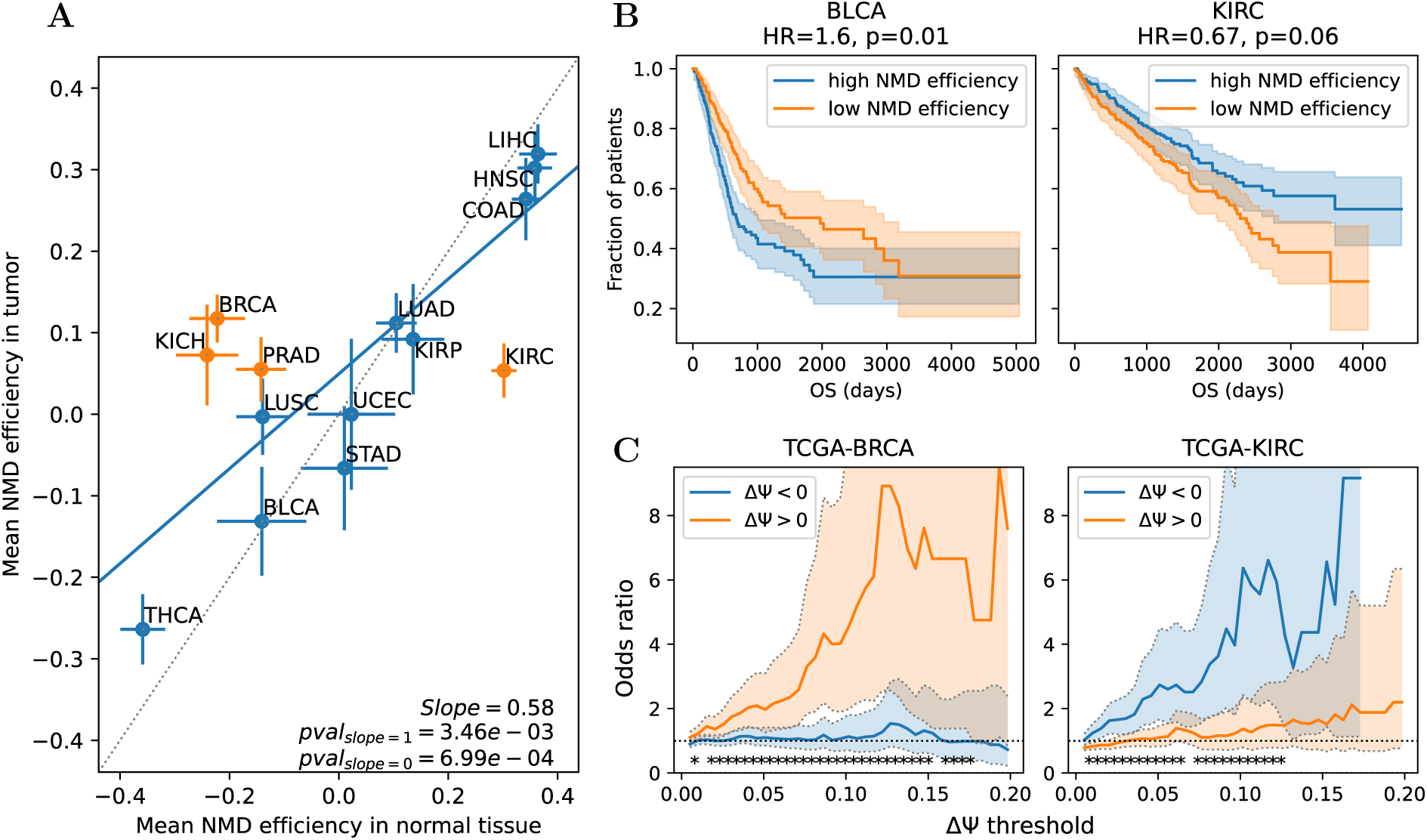
Association of NMD efficiency with patient survival and regulated unproductive splicing. **(A)** The least squares regression of the mean NMD efficiency in tumor versus that in normal tissue. Error bars represent standard errors of the mean. The dashed line corresponds to *y* = *x*. Tumors with a statistically significant difference in NMD efficiency are highlighted (*P* < 0.05, Wilcoxon test). **(B)** Kaplan-Meier curves for patient groups with high and low NMD efficiency in tumors, for bladder carcinoma (BLCA) and clear cell renal cell carcinoma (KIRC). **(C)** Odds ratio of oppositely directed changes in splicing and gene expression as a function of the threshold for the |ΔT| value and its sign. Asterisks denote a significant (*P* < 0.05) difference in odds ratios for positive and negative splicing changes.

We next assessed the clinical relevance of NMD efficiency alterations by relating it to patient survival. Significant associations with overall survival rate were observed in BLCA and KIRC (Figure 4B). Notably, the direction of these associations mirrored the baseline NMD efficiency levels of the normal tissue. In BLCA, which originally has low NMD efficiency, the elevated NMD efficiency in tumors correlated with poorer prognosis. Conversely, in KIRC, which exhibits high baseline NMD efficiency, the reduction of its activity was associated with unfavorable outcomes (Figure 4B). Together, these observations indicate that the greater the deviation of tumor NMD efficiency from the tissue-specific baseline, the worse the clinical prognosis, and this trend holds for both positive and negative deviations.

As we showed earlier, a substantial fraction of cancer-associated unproductive splicing changes can be attributed to global shifts in NMD efficiency, but some events may be regulated in a specific way. The event-specific regulation is expected to produce opposite changes in splicing and in the expression level of the host gene [42]. Using this negative association, we found that the fraction of specifically regulated USEs increased as we applied more stringent thresholds on the magnitude and and statistical significance of splicing changes (Figure S3). Moreover, these regulated USEs dominated among the events that shift against the global direction imposed by the changes in NMD efficiency. For example, in BRCA, the cancer type with the strongest increase in NMD efficiency, the regulated USEs were significantly enriched among positive splicing changes, whereas in KIRC, which exhibited the strongest decrease in NMD efficiency, the opposite pattern was observed (Figure 4C).

### The NMD efficiency metric recovers known and suggests novel regulators

The core components of the NMD pathway including the UPF proteins (UPF1, UPF2, UPF3), the SMG kinases and adaptor proteins (SMG1, SMG5, SMG6, SMG7), and the exon junction complex (EIF4A3, RBM8A, MAGOH) has been characterized in a considerable detail [43, 44, 45]. Still, several recent studies suggested that the regulatory landscape of NMD extends beyond these canonical components [46, 47]. To explore this broader scope, we applied the NMD efficiency metric to a compendium of 587 RBP knockdown and knockout experiments and asked which perturbations alter NMD globally [48, 28].

Most perturbations resulted in a reduced NMD efficiency, with exon junction complex components strongly being enriched among those with the largest effects (Figure 5A). Decreased NMD efficiency was also frequently observed upon depletion of spliceosomal factors, in line with observations implicating components of the spliceosome in the NMD control [47]. The pervasive tendency toward lower NMD efficiency upon depletion of almost any RBP likely reflects the high interconnectivity of the RBP regulatory network, in which perturbing a single RBP can cascade into global deregulation of RNA processing and accumulation of NMD-sensitive transcripts that are normally suppressed by both NMD and splicing regulatory programs [49].

**Figure 5:**
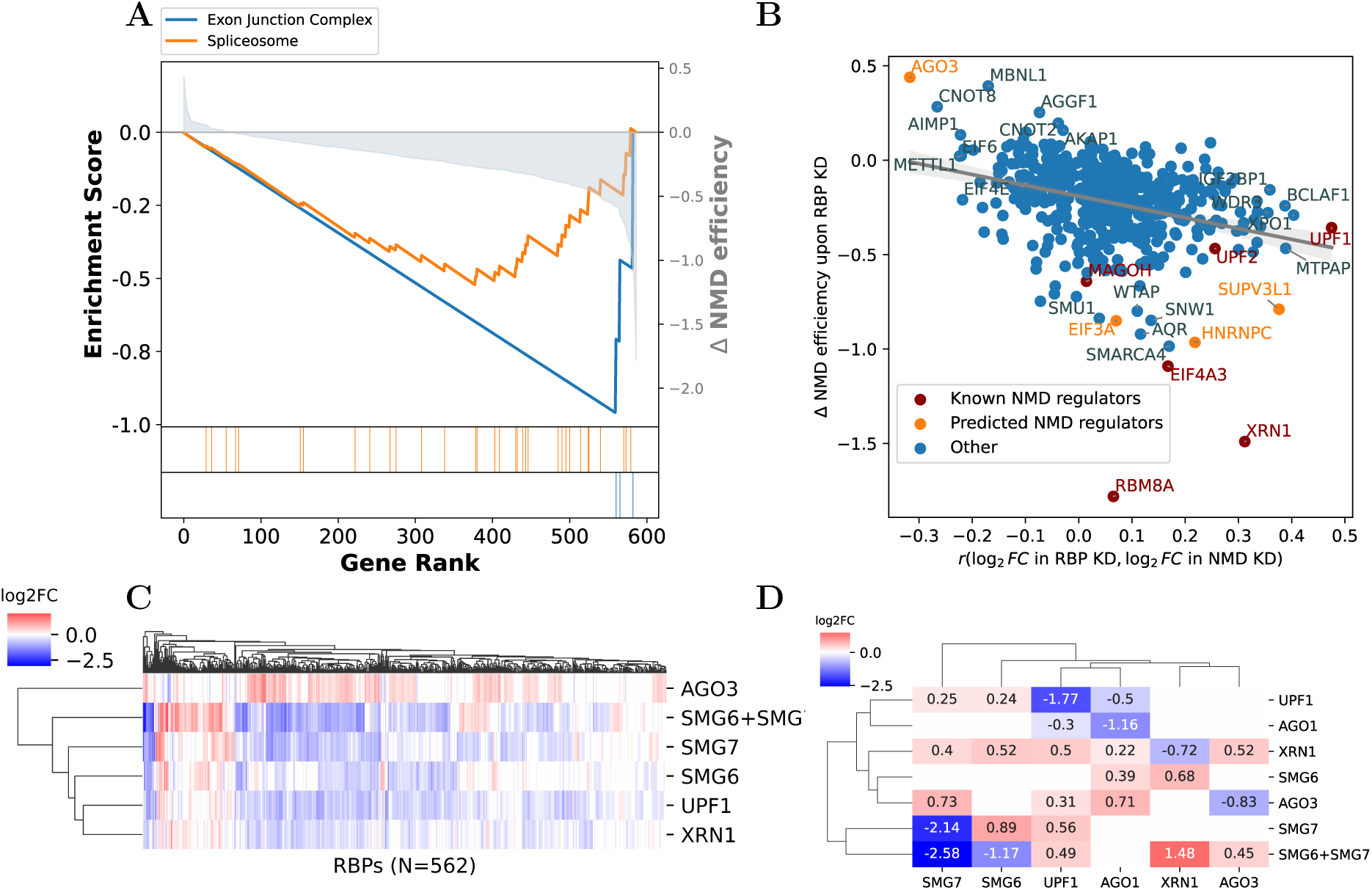
NMD efficiency in RBP inactivation experiments. **(A)** Gene set enrichment analysis (GSEA) plot for RBP ranked by the magnitude of NMD efficiency change in expression inactivation experiments. **(B)** Correlation between the change in NMD efficiency and the similarity of transcriptional profiles upon RBP depletion and NMD inactivation. The latter was assessed using the Pearson correlation coefficient applied to the log_2_ fold change of expression values (log_2_ *FC*) of differentially expressed genes in the RBP inactivation experiment and the NMD inactivation experiment (average over four NMD inactivation experiments). **(C)** Heat map of log_2_ *FC* for RBP genes upon knockdowns of AGO3 and NMD factors. **(D)** A part of the heat map from (B) that corresponds to expression changes of AGO1, AGO3, and NMD factors. Statistically insignificant changes are not shown.

To further characterize potential NMD regulators, we compared the gene expression changes that were induced by NMD inactivation with those observed after depletion of each RBP. Specifically, we related two quantities: (1) the change in NMD efficiency score under RBP depletion, and (2) the similarity of transcriptional profiles upon RBP depletion and NMD inactivation (the average of four NMD inactivation experiments: UPF1, SMG6, SMG7 and SMG6+SMG7). The latter was measured by the Pearson correlation coefficient of log_2_ fold change values of expression values of differentially expressed genes. RBPs whose inactivation reduces NMD efficiency tended to drive expression changes resembling those caused by NMD loss itself (Figure 5B). Among RBPs not previously linked to NMD, SUPV3L1, HNRNPC and EIF3A stood out as the closest to *bona fide* NMD factors both by the NMD efficiency metric and by their transcriptomic signatures.

SUPV3L1 is a helicase best known for its role in mitochondrial RNA turnover, although small amounts of this protein also localize to the nucleus [50]. Our data suggest that SUPV3L1 may also contribute to degradation of nuclear RNAs since the core USE events are mostly in nuclear genes, and they respond significantly to SUPV3L1 knockdown. HNRNPC binds poly- U tracts in mRNAs and is implicated in alternative splicing, polyadenylation, and translation [51, 52, 53, 54]. HNRNPC is known to suppress incorporation of numerous cryptic exons [55, 56], thus its impact on unproductive splicing may arise from event-specific regulation rather than direct involvement in NMD. EIF3A is the RNA-binding subunit of the eIF3 translation initiation complex, which was shown to interact physically with UPF1 and inhibit subsequent rounds of translation [57]. Knockdown of EIF3E, another eIF3 subunit, was previously shown to suppress NMD [58, 59]. Taken together, these observations point to reciprocal regulation between translation initiation and NMD.

We also identified putative negative regulators of NMD including AGO3, whose depletion produced the strongest increase in NMD efficiency across all tested RBPs. AGO3 is one of the four Argonaute genes involved in miRNA-guided translational repression [60]. Reporter assays mimicking RISC binding in 3’UTRs of NMD-sensitive transcripts previously showed that its paralog, AGO2, can inhibit NMD by suppressing translation of NMD-sensitive transcripts, although the extent to which this affects endogenous targets remains unclear [61]. Our data indicate that AGO3 suppresses degradation of a broad set of endogenous transcripts, leading to gene expression changes opposite to those caused by NMD inhibition (Figure 5C). However, other components of the miRNA pathway (AGO1, DICER1, DROSHA, XPO5, DGCR8) did not show similar effects, suggesting that the role of AGO3 is not simply a consequence of canonical miRNA pathway activity. Notably, AGO3 knockdown, but not AGO1 knockdown, increased expression of SMG7, which may partly account for the observed increase in NMD efficiency (Figure 5D). These findings point to AGO3 as a candidate negative regulator of NMD and suggest that it may have functions distinct from those of its paralogs.

## Discussion and conclusions

Accurate assessment of the NMD pathway efficiency is essential for understanding its contribution to cancer biology. The expression levels of NMD factors are unsuitable as proxies for pathway activity, as core NMD components are subject to negative autoregulation [21]. Existing approaches for estimating NMD efficiency include reporter assays (e.g., globin reporters) [25], analysis of allele-specific expression in genes with heterozygous nonsense mutations [23, 24], and comparison of expression levels of NMD-sensitive and productive transcripts [26]. Each method has its limitations: reporter assays are low throughput, allele-specific approaches rely on rare mutations that may be insufficiently represented in a sample [23, 26], and transcript-level quantification of NMD targets can be inaccurate for minor isoforms and sensitive to annotation errors [62].

In this study, we developed an NMD efficiency metric based on the splicing rates in a core set of USEs. We used it to characterize NMD regulation across human tissues, cancers, and large-scale RBP perturbation datasets. Leveraging USEs as endogenous reporters of NMD provides a powerful alternative to existing transcript- or mutation-based metrics. Using this framework, we confirmed that NMD efficiency varies in a non-random manner across human tissues, quantified the magnitude of these differences relative to NMD inhibition experiments, revealed systematic shifts of NMD activity in tumors, and identified putative novel regulators of the NMD pathway.

An unexpected observation emerging from our analyses and previous reports is the substantial heterogeneity of NMD efficiency within individual tissues [26]. The main factor contributing to this variability is that tissues in GTEx and TCGA represent mixtures of multiple cell types varying in their morphology, histology, and characteristic expression signatures [63]. Because NMD efficiency can vary between the individual cell types that make up a tissue [64], the NMD efficiency metric reflects not only the tissue-specific NMD system activity but also its histological composition. Overall, these results underscore the importance of considering tissue heterogeneity when interpreting any NMD efficiency estimates from RNA-seq.

Although tissue rankings by NMD efficiency produced by our metric and those from [23, 26] were consistent, our metric assigned higher NMD efficiency estimates for most brain regions (except Cerebellum, Anterior cingulate cortex (BA24), Hypothalamus and Spinal cord cervical C-1). Transcriptional programs of brain cells, particularly neurons, are known to be tightly regulated by unproductive splicing [65, 66, 67, 68], which may confound NMD efficiency estimates, if a substantial portion of core observations (USEs, mutations or transcript pairs) are specifically regulated. Indeed, we observed two clusters among core USEs, the larger one consisting of 91 USEs with lower T values in brain samples and the smaller, with 20 USEs having discordantly high T values in brain, yet being concordant with the majority of core USEs in samples from other tissues (Figure S4). Since our metric puts more strict requirements on the number of core USEs per sample than those of [23] and [26], with median 108 core USEs, compared to not more than 50 transcript pairs [26] and three stop gain variants [23] per sample in GTEx, it may provide a more realistic and conservative view on NMD efficiency in the brain. In multiple cancer types, we observed a striking divergence of NMD efficiency from the tissue-specific baseline. Tissues with intrinsically low NMD efficiency (e.g., breast) tend to exhibit an increase in tumors, whereas those with high baseline activity (e.g., kidney) often show reduced NMD activity. This “flattening” of NMD profiles suggests that tumors partially erase the NMD signature of their tissue of origin. One possible explanation is the acquisition of stem-like properties by cancer cells [69]. Stem and progenitor cells in many conditions display NMD efficiencies distinct from mature differentiated cells [70, 71, 4], and tumor dedifferentiation may shift NMD toward these stem-associated states. Alternatively, tumor evolution imposes selective pressures, such as chronic stress, altered translation demands, or immune evasion, that may favor specific levels of NMD independent of the original tissue program. Consistent with this, deviations from the tissue-specific NMD level were associated with patient survival, suggesting that maintenance of physiological NMD activity may be unfavorable for tumor progression.

Together with the observed pattern of co-directional changes of unproductive splicing, our results indicate a substantial degree of deregulation of the NMD pathway activity in tumors which undoubtedly contributes to the pathology of cancer.

## LIST OF ABBREVIATIONS

NMD: Nonsense mediated decay
USE: unproductive splicing event
RBP: RNA-binding protein
RNA-seq: RNA sequencing.

