## Supplementary material for "Global changes in unproductive splicing and the NMD system efficiency in tumors"

December 10, 2025

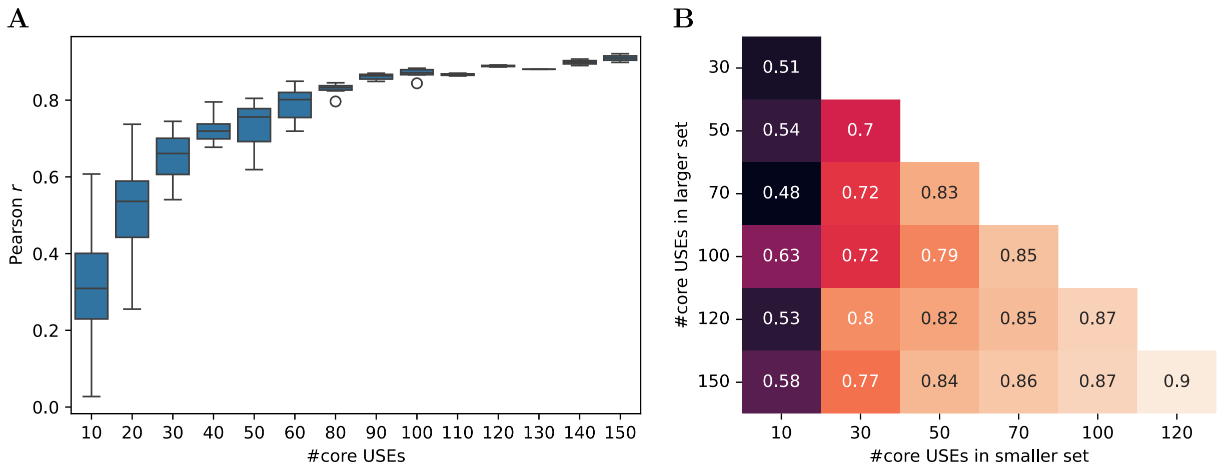


**Figure S1: (A)** Pearson correlation coefficients of NMD efficiencies in TCGA obtained from random USE sets as a function of their size. **(B)** Average pairwise Pearson correlation coefficients (5 replicates) of NMD efficiencies obtained from random USE sets of different sizes.

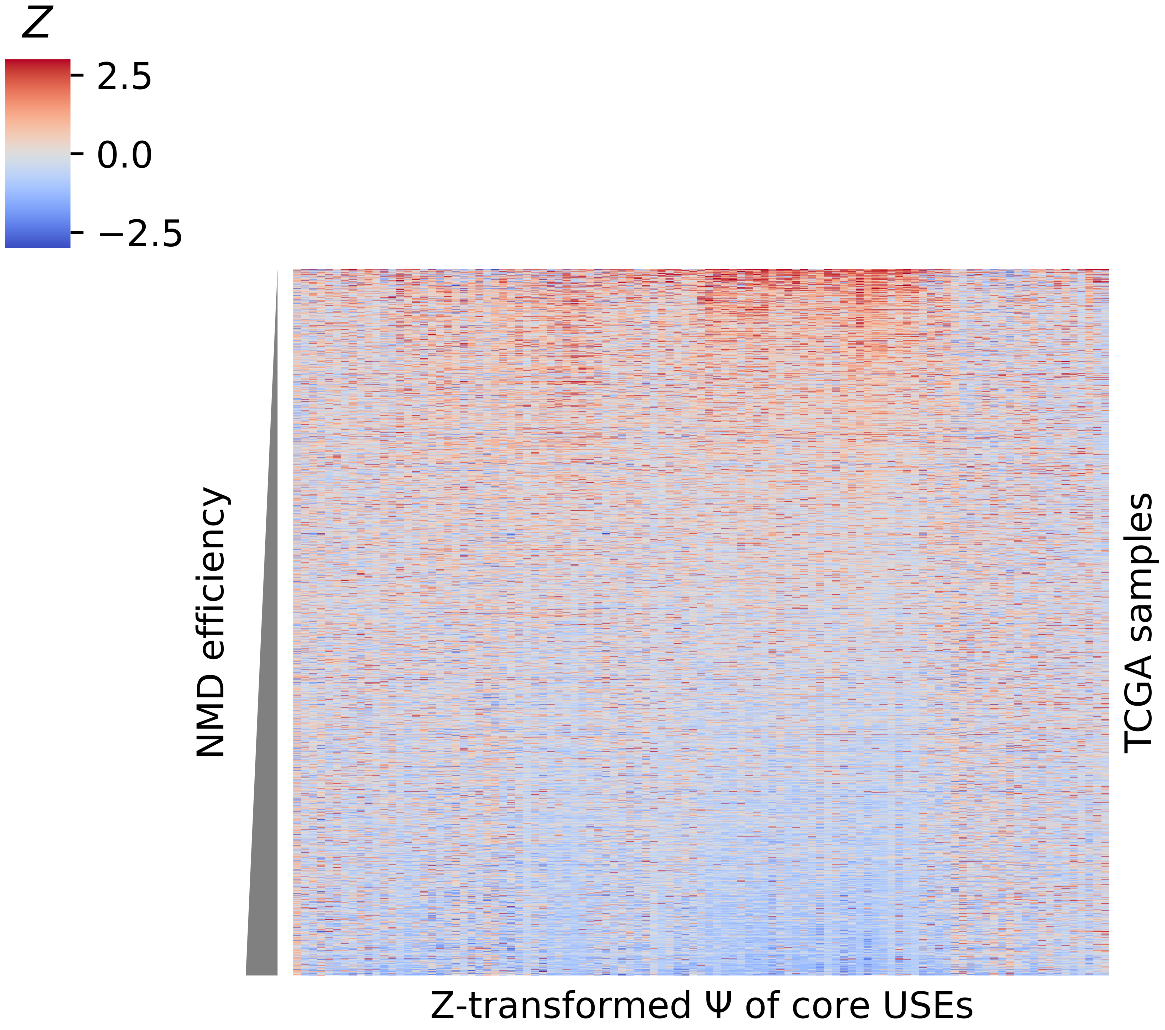


**Figure S2:** A heatmap of $z$-transformed $\Psi$ values of the core USE in TCGA samples. Samples are sorted by NMD efficiency that was estimated from $\Psi$ values.

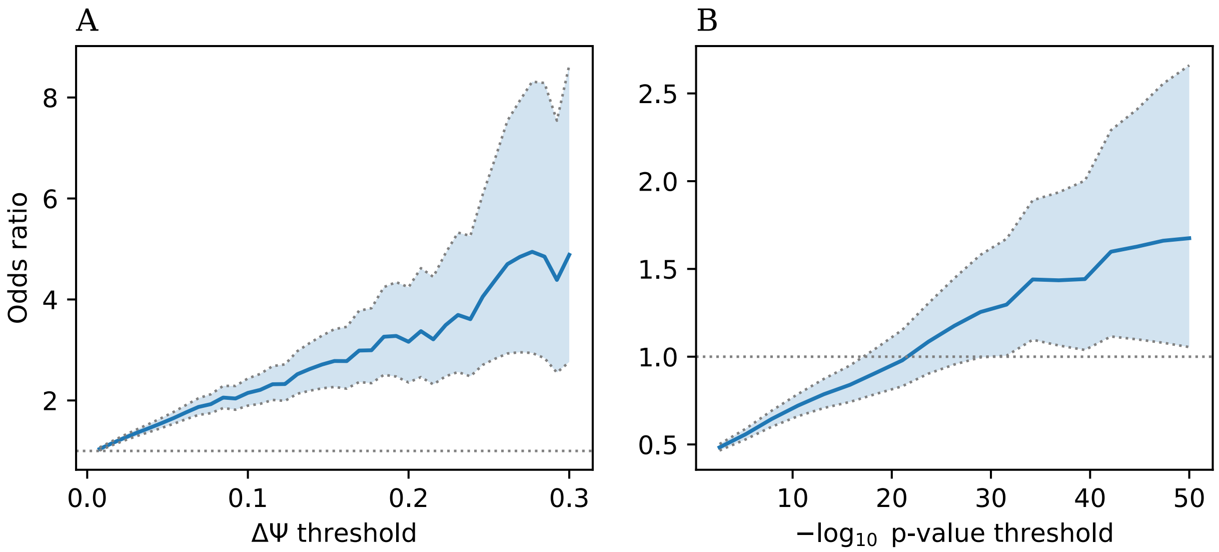


**Figure S3: (A, B)** Pan-cancer enrichment of specifically regulated USEs with increasing threshold on the magnitude of splicing change in the tumor (A) or on its statistical significance (B), calculated as the odds ratio of oppositely directed changes in splicing and gene expression among USEs with $\Delta\Psi$ values and $P$-values greater and less than a given threshold.

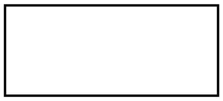

**Figure S4:** A heatmap of $z$-transformed $\Psi$ values of the core USEs in GTEx samples. Row colors indicate tissues (as in Fig. 3). The USE cluster with a specific splicing level in the brain is highlighted by a rectangle.

| TCGA cohort | Paired normal | Paired tumor | Normal | Tumor |
|---|---|---|---|---|
| TCGA-BLCA | 19 | 20 | 19 | 414 |
| TCGA-UCEC | 23 | 23 | 35 | 552 |
| TCGA-KICH | 23 | 23 | 24 | 65 |
| TCGA-STAD | 27 | 27 | 32 | 375 |
| TCGA-KIRP | 31 | 31 | 32 | 289 |
| TCGA-COAD | 41 | 44 | 41 | 480 |
| TCGA-HNSC | 43 | 42 | 44 | 501 |
| TCGA-LUSC | 49 | 49 | 49 | 502 |
| TCGA-LIHC | 50 | 50 | 50 | 374 |
| TCGA-PRAD | 52 | 54 | 52 | 500 |
| TCGA-LUAD | 57 | 63 | 59 | 536 |
| TCGA-THCA | 58 | 58 | 58 | 509 |
| TCGA-KIRC | 72 | 72 | 72 | 539 |
| TCGA-BRCA | 112 | 118 | 113 | 1108 |

**Table S1:** The number of tumor and matched normal tissue samples and number of paired tumor-normal samples in 13 TCGA cohorts that were selected for the analysis.

**Table S2:** The number of tissue samples in the GTEx dataset analyzed.

| Tissue | Number of samples |
|---|---|
| Adipose - Subcutaneous | 379 |
| Adipose - Visceral (Omentum) | 234 |
| Adrenal Gland | 159 |
| Artery - Aorta | 246 |
| Artery - Coronary | 140 |
| Artery - Tibial | 357 |
| Bladder | 10 |

**Table S2 – continued**

| | |
|---|---:|
| Brain - Amygdala | 81 |
| Brain - Anterior cingulate cortex (BA24) | 99 |
| Brain - Caudate (basal ganglia) | 134 |
| Brain - Cerebellar Hemisphere | 115 |
| Brain - Cerebellum | 144 |
| Brain - Cortex | 132 |
| Brain - Frontal Cortex (BA9) | 117 |
| Brain - Hippocampus | 103 |
| Brain - Hypothalamus | 104 |
| Brain - Nucleus accumbens (basal ganglia) | 123 |
| Brain - Putamen (basal ganglia) | 103 |
| Brain - Spinal cord (cervical c-1) | 76 |
| Brain - Substantia nigra | 71 |
| Breast - Mammary Tissue | 218 |
| Cells - Cultured fibroblasts | 305 |
| Cells - EBV-transformed lymphocytes | 131 |
| Cells - Leukemia cell line (CML) | 101 |
| Cervix - Ectocervix | 6 |
| Cervix - Endocervix | 5 |
| Colon - Sigmoid | 173 |
| Colon - Transverse | 202 |
| Esophagus - Gastroesophageal Junction | 176 |
| Esophagus - Mucosa | 330 |
| Esophagus - Muscularis | 283 |
| Fallopian Tube | 7 |
| Heart - Atrial Appendage | 218 |
| Heart - Left Ventricle | 267 |
| Kidney - Cortex | 36 |

**Table S2 – continued**

| | |
|---|---|
| Liver | 136 |
| Lung | 372 |
| Minor Salivary Gland | 70 |
| Muscle - Skeletal | 470 |
| Nerve - Tibial | 334 |
| Ovary | 108 |
| Pancreas | 192 |
| Pituitary | 124 |
| Prostate | 118 |
| Skin - Not Sun Exposed (Suprapubic) | 271 |
| Skin - Sun Exposed (Lower leg) | 392 |
| Small Intestine - Terminal Ileum | 104 |
| Spleen | 118 |
| Stomach | 203 |
| Testis | 198 |
| Thyroid | 354 |
| Uterus | 90 |
| Vagina | 97 |
| Whole Blood | 964 |

**Table S3:** List of RBP depletion experiments from ENCODE.

| RBP | Type | Cell line | Expt ID | Control ID |
|---|---|---|---|---|
| AARS | KD | HepG2 | ENCSR547NWD | ENCSR856ZRV |
| AARS | KD | K562 | ENCSR599UDS | ENCSR572FFX |
| AATF | KD | HepG2 | ENCSR424YSV | ENCSR003EKR |
| AATF | KD | K562 | ENCSR973QSV | ENCSR815CVQ |
| ABCF1 | KD | HepG2 | ENCSR610VTA | ENCSR067GHD |

**Table S3 – continued**

| | | | | |
|---|---|---|---|---|
| ABCF1 | KD | K562 | ENCSR721MXZ | ENCSR032YMP |
| ABT1 | KD | HepG2 | ENCSR756VLW | ENCSR225PRV |
| ABT1 | KD | K562 | ENCSR233UVM | ENCSR260BQC |
| ACO1 | KD | HepG2 | ENCSR511SYK | ENCSR538QOG |
| ACO1 | KO | K562 | ENCSR237QLO | ENCSR292PXV |
| ACO2 | KO | HepG2 | ENCSR251SXS | ENCSR953VMB |
| ACO2 | KO | K562 | ENCSR274ESZ | ENCSR900GQL |
| ADAR | KD | HepG2 | ENCSR104OLN | ENCSR305XWT |
| ADAR | KD | K562 | ENCSR164TLB | ENCSR667PLJ |
| ADAT1 | KO | HepG2 | ENCSR051LWN | ENCSR953VMB |
| ADAT1 | KO | K562 | ENCSR122RJE | ENCSR900GQL |
| ADAT3 | KO | HepG2 | ENCSR840HSL | ENCSR671FDZ |
| ADAT3 | KO | K562 | ENCSR994EWL | ENCSR877JEM |
| ADD3 | KO | HepG2 | ENCSR558DZW | ENCSR953VMB |
| ADK | KO | HepG2 | ENCSR924AVW | ENCSR953VMB |
| ADK | KO | K562 | ENCSR812JYP | ENCSR900GQL |
| AGFG1 | KO | HepG2 | ENCSR892ZGW | ENCSR953VMB |
| AGFG1 | KO | K562 | ENCSR654SOD | ENCSR900GQL |
| AGGF1 | KD | K562 | ENCSR812TLY | ENCSR143COQ |
| AGO1 | KD | HepG2 | ENCSR533HXS | ENCSR003EKR |
| AGO1 | KD | K562 | ENCSR268JDD | ENCSR084SCN |
| AGO2 | KD | K562 | ENCSR495YSS | ENCSR898NWE |
| AGO3 | KD | K562 | ENCSR207QGW | ENCSR898NWE |
| AHCYL2 | KO | HepG2 | ENCSR363NMO | ENCSR953VMB |
| AHCYL2 | KO | K562 | ENCSR700JXJ | ENCSR900GQL |
| AIMP1 | KO | HepG2 | ENCSR512BIT | ENCSR650TTU |
| AIMP1 | KO | K562 | ENCSR929YET | ENCSR900GQL |
| AKAP1 | KD | HepG2 | ENCSR016IDR | ENCSR237YZT |

**Table S3 – continued**

| | | | | |
|---|---|---|---|---|
| AKAP1 | KD | K562 | ENCSR338CON | ENCSR913CAE |
| AKAP8 | KD | HepG2 | ENCSR958KSY | ENCSR246RRQ |
| AKAP8 | KD | K562 | ENCSR000YYN | ENCSR143COQ |
| AKAP8L | KD | HepG2 | ENCSR807ODB | ENCSR538QOG |
| AKAP8L | KD | K562 | ENCSR809ISU | ENCSR143COQ |
| ALDH18A1 | KO | K562 | ENCSR642RCV | ENCSR900GQL |
| ALDOA | KO | HepG2 | ENCSR735FXX | ENCSR650TTU |
| ALDOA | KO | K562 | ENCSR071DBC | ENCSR900GQL |
| ALKBH1 | KO | HepG2 | ENCSR076LFC | ENCSR671FDZ |
| ALKBH1 | KO | K562 | ENCSR278CLE | ENCSR877JEM |
| APEH | KO | HepG2 | ENCSR828XJV | ENCSR650TTU |
| APEH | KO | K562 | ENCSR785JUR | ENCSR900GQL |
| APEX1 | KO | HepG2 | ENCSR367BWL | ENCSR750CPT |
| APEX1 | KO | K562 | ENCSR989ZNZ | ENCSR712CQC |
| API5 | KO | HepG2 | ENCSR585MSI | ENCSR650TTU |
| API5 | KO | K562 | ENCSR947EFJ | ENCSR900GQL |
| APOBEC3C | KD | HepG2 | ENCSR012DAF | ENCSR279HMU |
| APOBEC3C | KD | K562 | ENCSR963RLK | ENCSR620PUP |
| AQR | KD | HepG2 | ENCSR431SXF | ENCSR225PRV |
| AQR | KD | K562 | ENCSR624OUI | ENCSR718EWL |
| ARHGEF1 | KO | HepG2 | ENCSR250QFO | ENCSR650TTU |
| ARHGEF2 | KO | HepG2 | ENCSR191LWZ | ENCSR462MFN |
| ARHGEF2 | KO | K562 | ENCSR233MJI | ENCSR900GQL |
| ASCC1 | KD | HepG2 | ENCSR193FFA | ENCSR237YZT |
| ASCC1 | KD | K562 | ENCSR253DCB | ENCSR913CAE |
| ATP5A1 | KO | HepG2 | ENCSR340ZLQ | ENCSR462MFN |
| ATP5A1 | KO | K562 | ENCSR328DHD | ENCSR900GQL |
| ATP5C1 | KD | HepG2 | ENCSR713OLV | ENCSR067GHD |

**Table S3 – continued**

| | | | | |
|---|---|---|---|---|
| ATP5C1 | KD | K562 | ENCSR231DXJ | ENCSR143COQ |
| ATXN2L | KO | HepG2 | ENCSR234UAM | ENCSR248WTR |
| ATXN2L | KO | K562 | ENCSR585UBH | ENCSR900GQL |
| AUH | KD | HepG2 | ENCSR409CSO | ENCSR997HCQ |
| AUH | KD | K562 | ENCSR395FYF | ENCSR718EWL |
| BARD1 | KO | HepG2 | ENCSR104RTM | ENCSR953VMB |
| BARD1 | KO | K562 | ENCSR427KVG | ENCSR877JEM |
| BAZ1A | KO | HepG2 | ENCSR197XHV | ENCSR670KAS |
| BAZ1A | KO | K562 | ENCSR825EPB | ENCSR516TLX |
| BCCIP | KD | HepG2 | ENCSR570CWH | ENCSR856ZRV |
| BCCIP | KD | K562 | ENCSR606QIX | ENCSR245BNJ |
| BCLAF1 | KD | HepG2 | ENCSR481AYC | ENCSR856ZRV |
| BCLAF1 | KD | K562 | ENCSR410ZPU | ENCSR245BNJ |
| BMS1 | KO | K562 | ENCSR778XFE | ENCSR271ZAB |
| BOP1 | KD | HepG2 | ENCSR925SYZ | ENCSR264TUE |
| BOP1 | KD | K562 | ENCSR775TMW | ENCSR424QCW |
| BRD2 | KO | K562 | ENCSR046XQE | ENCSR900GQL |
| BRIX1 | KO | HepG2 | ENCSR060SLE | ENCSR671FDZ |
| BRIX1 | KO | K562 | ENCSR162HOU | ENCSR877JEM |
| BUD13 | KD | HepG2 | ENCSR382QKD | ENCSR067GHD |
| BUD13 | KD | K562 | ENCSR267RHP | ENCSR032YMP |
| BYSL | KO | K562 | ENCSR012OCK | ENCSR900GQL |
| BZW2 | KO | HepG2 | ENCSR921IQU | ENCSR650TTU |
| BZW2 | KO | K562 | ENCSR528OVC | ENCSR900GQL |
| C1QBP | KO | HepG2 | ENCSR944KLG | ENCSR650TTU |
| CACTIN | KO | HepG2 | ENCSR865WQR | ENCSR704VPL |
| CACTIN | KO | K562 | ENCSR416WTE | ENCSR877JEM |
| CALR | KD | HepG2 | ENCSR040WAK | ENCSR364GRM |

**Table S3 – continued**

| | | | | |
|---|---|---|---|---|
| CAND1 | KO | HepG2 | ENCSR297ONN | ENCSR650TTU |
| CAND1 | KO | K562 | ENCSR527XZI | ENCSR900GQL |
| CAPRIN1 | KO | K562 | ENCSR633RMN | ENCSR900GQL |
| CARHSP1 | KO | HepG2 | ENCSR974KDR | ENCSR650TTU |
| CARHSP1 | KO | K562 | ENCSR876IJP | ENCSR900GQL |
| CARS | KO | HepG2 | ENCSR077MFQ | ENCSR650TTU |
| CARS | KO | K562 | ENCSR267SUE | ENCSR900GQL |
| CAST | KO | K562 | ENCSR159HPD | ENCSR900GQL |
| CAT | KO | HepG2 | ENCSR051BQF | ENCSR650TTU |
| CAT | KO | K562 | ENCSR667JRK | ENCSR900GQL |
| CCAR1 | KD | HepG2 | ENCSR081IAO | ENCSR135LXL |
| CCAR1 | KD | K562 | ENCSR386YEV | ENCSR667PLJ |
| CCAR2 | KD | HepG2 | ENCSR237IWZ | ENCSR674KEK |
| CCAR2 | KD | K562 | ENCSR984CLJ | ENCSR620PUP |
| CCDC124 | KD | K562 | ENCSR874ZLI | ENCSR143COQ |
| CCDC47 | KO | HepG2 | ENCSR583SHC | ENCSR650TTU |
| CCNT1 | KO | K562 | ENCSR035MKJ | ENCSR900GQL |
| CCT2 | KO | K562 | ENCSR688ZJS | ENCSR220DTX |
| CCT3 | KO | K562 | ENCSR054DGN | ENCSR220DTX |
| CCT4 | KO | K562 | ENCSR995XEP | ENCSR220DTX |
| CD2BP2 | KO | K562 | ENCSR383FCI | ENCSR271ZAB |
| CD3EAP | KO | HepG2 | ENCSR400XYB | ENCSR964HKT |
| CD3EAP | KO | K562 | ENCSR323SDL | ENCSR220DTX |
| CDC40 | KO | HepG2 | ENCSR278NFF | ENCSR481WJH |
| CDC40 | KO | K562 | ENCSR354DVY | ENCSR015KGT |
| CDC42EP4 | KO | K562 | ENCSR323RFB | ENCSR271ZAB |
| CDV3 | KO | HepG2 | ENCSR875UMY | ENCSR462MFN |
| CDV3 | KO | K562 | ENCSR944PAO | ENCSR220DTX |

**Table S3 – continued**

| | | | | |
|---|---|---|---|---|
| CEBPZ | KD | HepG2 | ENCSR929PXS | ENCSR067GHD |
| CELF1 | KD | HepG2 | ENCSR695XOD | ENCSR042QTH |
| CELF1 | KD | K562 | ENCSR605MFS | ENCSR620PUP |
| CIRBP | KD | HepG2 | ENCSR230ORC | ENCSR237YZT |
| CIRBP | KD | K562 | ENCSR056QEW | ENCSR129RWD |
| CIRBP | KO | HepG2 | ENCSR010XIH | ENCSR827PII |
| CIRBP | KO | K562 | ENCSR452KAP | ENCSR292PXV |
| CKAP4 | KD | HepG2 | ENCSR269SJB | ENCSR538QOG |
| CLASP2 | KO | K562 | ENCSR331MLG | ENCSR220DTX |
| CLNS1A | KO | HepG2 | ENCSR513CVF | ENCSR462MFN |
| CLNS1A | KO | K562 | ENCSR950KYU | ENCSR220DTX |
| CLTC | KO | HepG2 | ENCSR576ASZ | ENCSR462MFN |
| CLTC | KO | K562 | ENCSR899RVY | ENCSR220DTX |
| CNDP2 | KO | HepG2 | ENCSR085GSF | ENCSR462MFN |
| CNDP2 | KO | K562 | ENCSR164POA | ENCSR220DTX |
| CNOT2 | KO | HepG2 | ENCSR918MUU | ENCSR671FDZ |
| CNOT2 | KO | K562 | ENCSR663BBL | ENCSR877JEM |
| CNOT7 | KD | HepG2 | ENCSR274KWA | ENCSR279HMU |
| CNOT7 | KD | K562 | ENCSR113PYX | ENCSR419JMU |
| CNOT8 | KD | K562 | ENCSR312HJY | ENCSR898NWE |
| CPEB4 | KD | K562 | ENCSR795VAK | ENCSR031RRO |
| CPSF2 | KO | HepG2 | ENCSR224XSS | ENCSR462MFN |
| CPSF2 | KO | K562 | ENCSR887XCL | ENCSR220DTX |
| CPSF3 | KO | HepG2 | ENCSR958OFR | ENCSR462MFN |
| CPSF3 | KO | K562 | ENCSR970WRV | ENCSR220DTX |
| CPSF4 | KO | HepG2 | ENCSR740WAG | ENCSR671FDZ |
| CPSF4 | KO | K562 | ENCSR854MSP | ENCSR854OFV |
| CPSF6 | KD | HepG2 | ENCSR676EKU | ENCSR264TUE |

**Table S3 – continued**

| | | | | |
|---|---|---|---|---|
| CPSF6 | KD | K562 | ENCSR384BDV | ENCSR424QCW |
| CPSF7 | KD | HepG2 | ENCSR594DNW | ENCSR067GHD |
| CPSF7 | KD | K562 | ENCSR222LRL | ENCSR667PLJ |
| CPSF7 | KO | HepG2 | ENCSR339XFB | ENCSR462MFN |
| CSDE1 | KO | HepG2 | ENCSR435YKS | ENCSR462MFN |
| CSE1L | KO | K562 | ENCSR877CJJ | ENCSR220DTX |
| CSNK1E | KO | K562 | ENCSR251LKQ | ENCSR220DTX |
| CSTF1 | KO | HepG2 | ENCSR751TUX | ENCSR704VPL |
| CSTF1 | KO | K562 | ENCSR529UYQ | ENCSR854OFV |
| CSTF2 | KD | HepG2 | ENCSR815JDY | ENCSR856ZRV |
| CSTF2 | KD | K562 | ENCSR885YOI | ENCSR245BNJ |
| CSTF2T | KD | HepG2 | ENCSR914WQV | ENCSR003EKR |
| CSTF2T | KD | K562 | ENCSR286OKW | ENCSR815CVQ |
| CSTF3 | KO | HepG2 | ENCSR044GQU | ENCSR462MFN |
| CTTN | KO | HepG2 | ENCSR755DOV | ENCSR462MFN |
| CTTN | KO | K562 | ENCSR146YRO | ENCSR220DTX |
| CWC15 | KO | K562 | ENCSR825RYR | ENCSR220DTX |
| CWF19L2 | KO | K562 | ENCSR582AOQ | ENCSR220DTX |
| DARS | KO | HepG2 | ENCSR574ZTO | ENCSR462MFN |
| DAZAP1 | KD | HepG2 | ENCSR220TBR | ENCSR042QTH |
| DAZAP1 | KD | K562 | ENCSR907UTB | ENCSR815CVQ |
| DBNL | KO | HepG2 | ENCSR256UIA | ENCSR462MFN |
| DBNL | KO | K562 | ENCSR593NFA | ENCSR220DTX |
| DCP1A | KO | K562 | ENCSR911ITT | ENCSR220DTX |
| DCP2 | KO | K562 | ENCSR423ERM | ENCSR854OFV |
| DCPS | KO | HepG2 | ENCSR524NOZ | ENCSR462MFN |
| DCPS | KO | K562 | ENCSR166VYI | ENCSR220DTX |
| DCTN2 | KO | HepG2 | ENCSR146ITQ | ENCSR462MFN |

**Table S3 – continued**

| | | | | |
|---|---|---|---|---|
| DDX1 | KD | HepG2 | ENCSR070LJO | ENCSR305XWT |
| DDX1 | KD | K562 | ENCSR208GPE | ENCSR667PLJ |
| DDX10 | KO | K562 | ENCSR002ODC | ENCSR220DTX |
| DDX18 | KO | HepG2 | ENCSR992YRL | ENCSR462MFN |
| DDX18 | KO | K562 | ENCSR300LXO | ENCSR220DTX |
| DDX19B | KD | HepG2 | ENCSR312SFA | ENCSR538QOG |
| DDX19B | KD | K562 | ENCSR281IUF | ENCSR143COQ |
| DDX21 | KD | HepG2 | ENCSR485ZTC | ENCSR003EKR |
| DDX21 | KD | K562 | ENCSR961WVL | ENCSR129RWD |
| DDX21 | KO | K562 | ENCSR384LKC | ENCSR163JUC |
| DDX23 | KO | K562 | ENCSR507WAD | ENCSR220DTX |
| DDX24 | KD | HepG2 | ENCSR300IEW | ENCSR003EKR |
| DDX24 | KD | K562 | ENCSR067LLB | ENCSR913CAE |
| DDX27 | KD | HepG2 | ENCSR210RWL | ENCSR856ZRV |
| DDX27 | KD | K562 | ENCSR584LDM | ENCSR245BNJ |
| DDX28 | KD | HepG2 | ENCSR222CSF | ENCSR856ZRV |
| DDX28 | KD | K562 | ENCSR205VSQ | ENCSR661HEL |
| DDX39B | KO | HepG2 | ENCSR175OSV | ENCSR670KAS |
| DDX39B | KO | K562 | ENCSR883MTV | ENCSR516TLX |
| DDX3X | KD | HepG2 | ENCSR637JLM | ENCSR237YZT |
| DDX3X | KD | K562 | ENCSR000KYM | ENCSR913CAE |
| DDX42 | KO | HepG2 | ENCSR718RBZ | ENCSR834IFU |
| DDX42 | KO | K562 | ENCSR521WCW | ENCSR015KGT |
| DDX43 | KO | K562 | ENCSR390TOZ | ENCSR877JEM |
| DDX47 | KD | HepG2 | ENCSR388CNS | ENCSR968BBQ |
| DDX47 | KD | K562 | ENCSR155EZL | ENCSR031RRO |
| DDX5 | KD | HepG2 | ENCSR808FBR | ENCSR776SXA |
| DDX51 | KD | K562 | ENCSR029LGJ | ENCSR661HEL |

**Table S3 – continued**

| | | | | |
|---|---|---|---|---|
| DDX51 | KO | HepG2 | ENCSR080NRY | ENCSR827PII |
| DDX52 | KD | HepG2 | ENCSR913ZWR | ENCSR689PHN |
| DDX52 | KD | K562 | ENCSR560RSZ | ENCSR661HEL |
| DDX55 | KD | HepG2 | ENCSR964YTW | ENCSR856ZRV |
| DDX55 | KD | K562 | ENCSR856CJK | ENCSR572FFX |
| DDX59 | KD | HepG2 | ENCSR598GKQ | ENCSR264TUE |
| DDX6 | KD | HepG2 | ENCSR147ZBD | ENCSR237YZT |
| DDX6 | KD | K562 | ENCSR119QWQ | ENCSR913CAE |
| DDX6 | KO | K562 | ENCSR298ASI | ENCSR015KGT |
| DGCR8 | KO | K562 | ENCSR148UGK | ENCSR619GAS |
| DHX30 | KD | HepG2 | ENCSR853PBF | ENCSR305XWT |
| DHX30 | KD | K562 | ENCSR345VVZ | ENCSR667PLJ |
| DHX57 | KO | HepG2 | ENCSR029VGU | ENCSR670KAS |
| DHX57 | KO | K562 | ENCSR359ICP | ENCSR516TLX |
| DICER1 | KO | HepG2 | ENCSR428WVZ | ENCSR964HKT |
| DICER1 | KO | K562 | ENCSR715XCS | ENCSR404OHQ |
| DKC1 | KD | HepG2 | ENCSR118KUN | ENCSR538QOG |
| DKC1 | KD | K562 | ENCSR494UDF | ENCSR143COQ |
| DNAJC2 | KD | HepG2 | ENCSR004OSI | ENCSR305XWT |
| DNAJC2 | KD | K562 | ENCSR577OVP | ENCSR667PLJ |
| DNAJC21 | KD | HepG2 | ENCSR385KOY | ENCSR305XWT |
| DNAJC21 | KD | K562 | ENCSR079LMZ | ENCSR667PLJ |
| DROSHA | KD | K562 | ENCSR624XHG | ENCSR164MUK |
| DROSHA | KO | HepG2 | ENCSR991XFO | ENCSR481WJH |
| DUS3L | KO | HepG2 | ENCSR020EBN | ENCSR670KAS |
| DUS3L | KO | K562 | ENCSR949QWR | ENCSR516TLX |
| EEF2 | KD | HepG2 | ENCSR477TRX | ENCSR689PHN |
| EEF2 | KD | K562 | ENCSR181RLB | ENCSR661HEL |

**Table S3 – continued**

| | | | | |
|---|---|---|---|---|
| EFTUD2 | KD | HepG2 | ENCSR620OKS | ENCSR689PHN |
| EFTUD2 | KD | K562 | ENCSR117WLY | ENCSR661HEL |
| EIF2AK1 | KO | K562 | ENCSR667WZB | ENCSR404OHQ |
| EIF2AK2 | KO | K562 | ENCSR628AFF | ENCSR712CQC |
| EIF2B3 | KO | HepG2 | ENCSR312UYN | ENCSR964HKT |
| EIF2B3 | KO | K562 | ENCSR786OLN | ENCSR404OHQ |
| EIF2S1 | KD | HepG2 | ENCSR861ENA | ENCSR237YZT |
| EIF2S1 | KD | K562 | ENCSR546MBH | ENCSR913CAE |
| EIF2S2 | KD | HepG2 | ENCSR110HAA | ENCSR067GHD |
| EIF2S2 | KD | K562 | ENCSR076PMZ | ENCSR118EFE |
| EIF3A | KD | K562 | ENCSR258VGD | ENCSR667PLJ |
| EIF3D | KD | HepG2 | ENCSR788HVK | ENCSR689PHN |
| EIF3D | KD | K562 | ENCSR660ETT | ENCSR661HEL |
| EIF3G | KD | HepG2 | ENCSR778AJO | ENCSR003EKR |
| EIF3G | KD | K562 | ENCSR143UET | ENCSR084SCN |
| EIF3H | KO | HepG2 | ENCSR733MVT | ENCSR805TIZ |
| EIF3H | KO | K562 | ENCSR096WVE | ENCSR226KWO |
| EIF4A3 | KD | HepG2 | ENCSR957EEG | ENCSR237YZT |
| EIF4A3 | KD | K562 | ENCSR961YAG | ENCSR913CAE |
| EIF4A3 | KO | HepG2 | ENCSR999MAE | ENCSR964HKT |
| EIF4A3 | KO | K562 | ENCSR264MSX | ENCSR404OHQ |
| EIF4B | KD | HepG2 | ENCSR313CHR | ENCSR237YZT |
| EIF4B | KD | K562 | ENCSR774BXV | ENCSR913CAE |
| EIF4B | KO | HepG2 | ENCSR323OPW | ENCSR775ZWO |
| EIF4E | KO | HepG2 | ENCSR294DOR | ENCSR964HKT |
| EIF4E | KO | K562 | ENCSR033XYV | ENCSR404OHQ |
| EIF4G1 | KD | HepG2 | ENCSR509LIV | ENCSR042QTH |
| EIF4G1 | KD | K562 | ENCSR712CSN | ENCSR667PLJ |

**Table S3 – continued**

| | | | | |
|---|---|---|---|---|
| EIF4G2 | KD | HepG2 | ENCSR152MON | ENCSR042QTH |
| EIF4G2 | KD | K562 | ENCSR040FSN | ENCSR620PUP |
| EIF6 | KO | HepG2 | ENCSR608WXK | ENCSR671FDZ |
| EIF6 | KO | K562 | ENCSR975NHV | ENCSR877JEM |
| ELAC2 | KO | HepG2 | ENCSR511EKJ | ENCSR671FDZ |
| ELAC2 | KO | K562 | ENCSR715GSX | ENCSR877JEM |
| ELAVL1 | KO | HepG2 | ENCSR659DRE | ENCSR750CPT |
| ELAVL1 | KO | K562 | ENCSR356PRP | ENCSR712CQC |
| ERO1A | KO | HepG2 | ENCSR561IMV | ENCSR964HKT |
| ERO1A | KO | K562 | ENCSR353OQP | ENCSR404OHQ |
| ESF1 | KD | HepG2 | ENCSR060IWW | ENCSR067GHD |
| ESF1 | KD | K562 | ENCSR077BPR | ENCSR032YMP |
| ETF1 | KD | HepG2 | ENCSR840QOH | ENCSR997HCQ |
| ETF1 | KO | K562 | ENCSR066ZMF | ENCSR015KGT |
| EWSR1 | KD | HepG2 | ENCSR532ZPP | ENCSR538QOG |
| EWSR1 | KD | K562 | ENCSR831YGP | ENCSR913CAE |
| EXOSC10 | KO | HepG2 | ENCSR455KXW | ENCSR704VPL |
| EXOSC10 | KO | K562 | ENCSR299PRA | ENCSR854OFV |
| EXOSC2 | KO | HepG2 | ENCSR035LEO | ENCSR704VPL |
| EXOSC2 | KO | K562 | ENCSR473HXR | ENCSR854OFV |
| EXOSC5 | KO | HepG2 | ENCSR284SZO | ENCSR521WAI |
| EXOSC5 | KO | K562 | ENCSR406SEH | ENCSR341TTW |
| EXOSC9 | KD | HepG2 | ENCSR597IYB | ENCSR003EKR |
| EXOSC9 | KD | K562 | ENCSR812EIA | ENCSR815CVQ |
| FAM120A | KD | HepG2 | ENCSR047VPW | ENCSR689PHN |
| FAM120A | KD | K562 | ENCSR492BKM | ENCSR661HEL |
| FASTKD1 | KD | HepG2 | ENCSR728BOL | ENCSR279HMU |
| FASTKD2 | KD | HepG2 | ENCSR716WZH | ENCSR003EKR |

**Table S3 – continued**

| | | | | |
|---|---|---|---|---|
| FASTKD2 | KD | K562 | ENCSR608IAI | ENCSR084SCN |
| FIP1L1 | KD | HepG2 | ENCSR116QBU | ENCSR067GHD |
| FIP1L1 | KD | K562 | ENCSR511BNY | ENCSR032YMP |
| FKBP4 | KD | HepG2 | ENCSR639LKS | ENCSR856ZRV |
| FKBP4 | KD | K562 | ENCSR379VXW | ENCSR572FFX |
| FMR1 | KD | HepG2 | ENCSR905HID | ENCSR042QTH |
| FMR1 | KD | K562 | ENCSR555LCE | ENCSR424QCW |
| FTO | KD | HepG2 | ENCSR389HFU | ENCSR264TUE |
| FTO | KD | K562 | ENCSR688GVV | ENCSR424QCW |
| FUBP1 | KD | HepG2 | ENCSR736TAW | ENCSR225PRV |
| FUBP1 | KD | K562 | ENCSR608IXR | ENCSR260BQC |
| FUBP3 | KD | HepG2 | ENCSR755KOM | ENCSR376RJN |
| FUBP3 | KD | K562 | ENCSR373KOF | ENCSR913CAE |
| FUS | KD | HepG2 | ENCSR927JXU | ENCSR491FOC |
| FUS | KD | K562 | ENCSR325OOM | ENCSR084SCN |
| FXR1 | KD | HepG2 | ENCSR009PPI | ENCSR237YZT |
| FXR1 | KD | K562 | ENCSR780YFF | ENCSR913CAE |
| FXR2 | KD | K562 | ENCSR139BIJ | ENCSR081EST |
| FXR2 | KO | HepG2 | ENCSR147RTA | ENCSR834IFU |
| FXR2 | KO | K562 | ENCSR376GHG | ENCSR828YEB |
| FYTTD1 | KO | K562 | ENCSR832XLB | ENCSR854OFV |
| G3BP1 | KD | HepG2 | ENCSR074UZM | ENCSR237YZT |
| G3BP1 | KD | K562 | ENCSR792CBM | ENCSR129RWD |
| G3BP1 | KO | K562 | ENCSR846OQD | ENCSR292PXV |
| G3BP2 | KD | HepG2 | ENCSR945UYL | ENCSR237YZT |
| G3BP2 | KD | K562 | ENCSR246SOU | ENCSR344XID |
| GARS | KO | HepG2 | ENCSR474EJY | ENCSR670KAS |
| GARS | KO | K562 | ENCSR503VIZ | ENCSR516TLX |

**Table S3 – continued**

| | | | | |
|---|---|---|---|---|
| GEMIN5 | KD | HepG2 | ENCSR771QMJ | ENCSR689PHN |
| GEMIN5 | KD | K562 | ENCSR398GHW | ENCSR661HEL |
| GLRX3 | KD | K562 | ENCSR874DVZ | ENCSR174OYC |
| GLRX3 | KO | HepG2 | ENCSR400SCX | ENCSR827PII |
| GNB2L1 | KD | HepG2 | ENCSR116YMU | ENCSR674KEK |
| GNL3 | KO | HepG2 | ENCSR940FIN | ENCSR713OTF |
| GNL3 | KO | K562 | ENCSR455VFL | ENCSR015KGT |
| GPKOW | KD | HepG2 | ENCSR968YWY | ENCSR264TUE |
| GPKOW | KD | K562 | ENCSR967QNT | ENCSR424QCW |
| GRSF1 | KD | HepG2 | ENCSR674KDQ | ENCSR279HMU |
| GRSF1 | KD | K562 | ENCSR835RMN | ENCSR118EFE |
| GRWD1 | KD | HepG2 | ENCSR850FEH | ENCSR674KEK |
| GRWD1 | KD | K562 | ENCSR528ASX | ENCSR572FFX |
| GSK3A | KO | HepG2 | ENCSR300RMY | ENCSR750CPT |
| GSK3A | KO | K562 | ENCSR610HJO | ENCSR712CQC |
| GSK3B | KO | HepG2 | ENCSR426JCD | ENCSR964HKT |
| GSK3B | KO | K562 | ENCSR025XBH | ENCSR404OHQ |
| GSPT2 | KO | K562 | ENCSR148XGX | ENCSR516TLX |
| GTF2F1 | KD | HepG2 | ENCSR295XKC | ENCSR491FOC |
| GTF2F1 | KD | K562 | ENCSR188IPO | ENCSR815CVQ |
| HDGF | KD | HepG2 | ENCSR634KHL | ENCSR538QOG |
| HDGF | KD | K562 | ENCSR793QPX | ENCSR619PCR |
| HLTF | KD | HepG2 | ENCSR010ZMZ | ENCSR491FOC |
| HLTF | KD | K562 | ENCSR958NDU | ENCSR815CVQ |
| HNRNPA0 | KD | HepG2 | ENCSR720BPO | ENCSR279HMU |
| HNRNPA0 | KD | K562 | ENCSR552NBS | ENCSR419JMU |
| HNRNPA1 | KD | HepG2 | ENCSR182DAW | ENCSR305XWT |
| HNRNPA1 | KD | K562 | ENCSR048BWH | ENCSR438MDN |

**Table S3 – continued**

| | | | | |
|---|---|---|---|---|
| HNRNPA2B1 | KD | HepG2 | ENCSR769GES | ENCSR042QTH |
| HNRNPA2B1 | KD | K562 | ENCSR794NUE | ENCSR164MUK |
| HNRNPAB | KD | HepG2 | ENCSR354XQY | ENCSR279HMU |
| HNRNPAB | KD | K562 | ENCSR778WPL | ENCSR419JMU |
| HNRNPC | KD | HepG2 | ENCSR052IYH | ENCSR305XWT |
| HNRNPC | KD | K562 | ENCSR634KBO | ENCSR572FFX |
| HNRNPD | KD | HepG2 | ENCSR660MZN | ENCSR279HMU |
| HNRNPF | KD | HepG2 | ENCSR693MZJ | ENCSR305XWT |
| HNRNPF | KD | K562 | ENCSR392HSJ | ENCSR572FFX |
| HNRNPF | KO | HepG2 | ENCSR599NNK | ENCSR105NML |
| HNRNPH1 | KO | HepG2 | ENCSR094HEU | ENCSR194SPW |
| HNRNPH1 | KO | K562 | ENCSR354RSR | ENCSR341TTW |
| HNRNPK | KD | HepG2 | ENCSR853ZJS | ENCSR237YZT |
| HNRNPK | KD | K562 | ENCSR529JNJ | ENCSR129RWD |
| HNRNPL | KD | HepG2 | ENCSR155BMF | ENCSR042QTH |
| HNRNPL | KD | K562 | ENCSR563YIS | ENCSR031RRO |
| HNRNPLL | KD | HepG2 | ENCSR490DYI | ENCSR264TUE |
| HNRNPM | KD | HepG2 | ENCSR995JMS | ENCSR067GHD |
| HNRNPM | KD | K562 | ENCSR746NIM | ENCSR032YMP |
| HNRNPU | KD | HepG2 | ENCSR308IKH | ENCSR305XWT |
| HNRNPU | KD | K562 | ENCSR047IUS | ENCSR438MDN |
| HNRNPUL1 | KD | HepG2 | ENCSR689ZJC | ENCSR104ABF |
| HNRNPUL1 | KD | K562 | ENCSR034VBA | ENCSR164MUK |
| HSPD1 | KD | HepG2 | ENCSR243IGA | ENCSR246RRQ |
| HSPD1 | KD | K562 | ENCSR222ABK | ENCSR032YMP |
| IFIT2 | KO | HepG2 | ENCSR144HCM | ENCSR670KAS |
| IFIT2 | KO | K562 | ENCSR671OGO | ENCSR516TLX |
| IGF2BP1 | KD | HepG2 | ENCSR708GKW | ENCSR491FOC |

**Table S3 – continued**

| | | | | |
|---|---|---|---|---|
| IGF2BP1 | KD | K562 | ENCSR629EWX | ENCSR129RWD |
| IGF2BP1 | KO | HepG2 | ENCSR268XHM | ENCSR105NML |
| IGF2BP1 | KO | K562 | ENCSR543LCG | ENCSR163JUC |
| IGF2BP2 | KD | HepG2 | ENCSR478FJK | ENCSR237YZT |
| IGF2BP2 | KD | K562 | ENCSR952RRH | ENCSR129RWD |
| IGF2BP3 | KD | HepG2 | ENCSR710NWE | ENCSR237YZT |
| IGF2BP3 | KD | K562 | ENCSR302JQA | ENCSR129RWD |
| ILF2 | KD | HepG2 | ENCSR366FFV | ENCSR305XWT |
| ILF2 | KD | K562 | ENCSR126ARZ | ENCSR419JMU |
| ILF3 | KD | HepG2 | ENCSR942MBU | ENCSR279HMU |
| ILF3 | KD | K562 | ENCSR269HQA | ENCSR667PLJ |
| KHDRBS1 | KD | HepG2 | ENCSR784FTX | ENCSR279HMU |
| KHDRBS1 | KD | K562 | ENCSR023HWI | ENCSR667PLJ |
| KHSRP | KD | HepG2 | ENCSR850CKU | ENCSR237YZT |
| KHSRP | KD | K562 | ENCSR561CBC | ENCSR129RWD |
| KIF1C | KD | HepG2 | ENCSR182GKG | ENCSR538QOG |
| KIF1C | KD | K562 | ENCSR823WTA | ENCSR143COQ |
| KPNB1 | KO | HepG2 | ENCSR977NJB | ENCSR750CPT |
| KPNB1 | KO | K562 | ENCSR667FZF | ENCSR712CQC |
| KRR1 | KD | HepG2 | ENCSR542ESY | ENCSR246RRQ |
| KRR1 | KD | K562 | ENCSR244SIO | ENCSR424QCW |
| LARP4 | KD | HepG2 | ENCSR744PAQ | ENCSR135LXL |
| LARP4 | KD | K562 | ENCSR866XLI | ENCSR164MUK |
| LARP7 | KD | HepG2 | ENCSR624FBY | ENCSR674KEK |
| LARP7 | KD | K562 | ENCSR770OWW | ENCSR572FFX |
| LARS | KO | HepG2 | ENCSR213ILK | ENCSR671FDZ |
| LARS | KO | K562 | ENCSR381NPB | ENCSR877JEM |
| LIN28B | KD | HepG2 | ENCSR927SLP | ENCSR674KEK |

**Table S3 – continued**

| | | | | |
|---|---|---|---|---|
| LIN28B | KD | K562 | ENCSR598YQX | ENCSR572FFX |
| LSM11 | KD | HepG2 | ENCSR883BXR | ENCSR674KEK |
| LSM11 | KD | K562 | ENCSR762FEO | ENCSR572FFX |
| LSM7 | KO | HepG2 | ENCSR120WNJ | ENCSR704VPL |
| LSM7 | KO | K562 | ENCSR307KLI | ENCSR854OFV |
| MAGOH | KD | HepG2 | ENCSR746EKS | ENCSR237YZT |
| MAGOH | KD | K562 | ENCSR849STR | ENCSR129RWD |
| MAK16 | KD | K562 | ENCSR517JHY | ENCSR143COQ |
| MAK16 | KO | HepG2 | ENCSR624YDI | ENCSR827PII |
| MARK2 | KD | HepG2 | ENCSR105OXX | ENCSR674KEK |
| MARK2 | KD | K562 | ENCSR016OIX | ENCSR619PCR |
| MATR3 | KD | HepG2 | ENCSR492UFS | ENCSR237YZT |
| MATR3 | KD | K562 | ENCSR792XFP | ENCSR129RWD |
| MBNL1 | KD | K562 | ENCSR222COT | ENCSR143COQ |
| MDM2 | KO | K562 | ENCSR943CXG | ENCSR978MMZ |
| METAP2 | KD | HepG2 | ENCSR992JGE | ENCSR689PHN |
| METAP2 | KD | K562 | ENCSR952QDQ | ENCSR424QCW |
| METTL1 | KO | HepG2 | ENCSR925POR | ENCSR671FDZ |
| METTL1 | KO | K562 | ENCSR607SRH | ENCSR877JEM |
| METTL3 | KO | HepG2 | ENCSR884REL | ENCSR704VPL |
| METTL3 | KO | K562 | ENCSR335KFN | ENCSR877JEM |
| MORC2 | KO | HepG2 | ENCSR444IOJ | ENCSR306EWE |
| MORC2 | KO | K562 | ENCSR990DKW | ENCSR978MMZ |
| MSI2 | KD | HepG2 | ENCSR896MMU | ENCSR067GHD |
| MSI2 | KD | K562 | ENCSR169QQW | ENCSR620PUP |
| MTPAP | KD | HepG2 | ENCSR701GSV | ENCSR264TUE |
| MTPAP | KD | K562 | ENCSR631RFX | ENCSR174OYC |
| NAA15 | KD | HepG2 | ENCSR355OQC | ENCSR067GHD |

**Table S3 – continued**

| | | | | |
|---|---|---|---|---|
| NAA15 | KD | K562 | ENCSR945GUR | ENCSR143COQ |
| NCBP1 | KO | HepG2 | ENCSR324GSI | ENCSR775ZWO |
| NCBP1 | KO | K562 | ENCSR090SBZ | ENCSR292PXV |
| NCBP2 | KD | HepG2 | ENCSR030ARO | ENCSR237YZT |
| NCBP2 | KD | K562 | ENCSR361LBE | ENCSR913CAE |
| NCL | KO | HepG2 | ENCSR013PBS | ENCSR670KAS |
| NCL | KO | K562 | ENCSR665CVU | ENCSR978MMZ |
| NELFE | KD | HepG2 | ENCSR939ZRA | ENCSR603TCV |
| NELFE | KD | K562 | ENCSR201WFU | ENCSR129RWD |
| NFX1 | KD | K562 | ENCSR007XKL | ENCSR815CVQ |
| NIP7 | KD | HepG2 | ENCSR696LLZ | ENCSR674KEK |
| NIP7 | KO | HepG2 | ENCSR586YGD | ENCSR300EBW |
| NIP7 | KO | K562 | ENCSR499IQB | ENCSR292PXV |
| NKRF | KD | HepG2 | ENCSR517JDK | ENCSR246RRQ |
| NKRF | KD | K562 | ENCSR231PWH | ENCSR619PCR |
| NOL12 | KD | HepG2 | ENCSR643UFV | ENCSR042QTH |
| NOL12 | KD | K562 | ENCSR227AVS | ENCSR620PUP |
| NOLC1 | KO | HepG2 | ENCSR959HPB | ENCSR521WAI |
| NOLC1 | KO | K562 | ENCSR410USU | ENCSR015KGT |
| NONO | KD | HepG2 | ENCSR647NYX | ENCSR856ZRV |
| NONO | KD | K562 | ENCSR398HXV | ENCSR661HEL |
| NPM1 | KD | HepG2 | ENCSR016XPB | ENCSR104ABF |
| NPM1 | KD | K562 | ENCSR346DZQ | ENCSR164MUK |
| NSUN2 | KD | HepG2 | ENCSR629RUG | ENCSR264TUE |
| NSUN4 | KO | HepG2 | ENCSR504CPZ | ENCSR964HKT |
| NSUN4 | KO | K562 | ENCSR550VEL | ENCSR404OHQ |
| NSUN5 | KO | K562 | ENCSR982XTO | ENCSR978MMZ |
| NUFIP2 | KD | HepG2 | ENCSR584JRB | ENCSR104ABF |

**Table S3 – continued**

| | | | | |
|---|---|---|---|---|
| NUFIP2 | KD | K562 | ENCSR754RJA | ENCSR174OYC |
| NUP35 | KD | HepG2 | ENCSR457WBK | ENCSR960MSV |
| NUP35 | KD | K562 | ENCSR953IQF | ENCSR260BQC |
| NUSAP1 | KD | HepG2 | ENCSR180XTP | ENCSR674KEK |
| NUSAP1 | KD | K562 | ENCSR927XBT | ENCSR572FFX |
| PA2G4 | KD | HepG2 | ENCSR028YAQ | ENCSR689PHN |
| PA2G4 | KD | K562 | ENCSR309PPC | ENCSR661HEL |
| PABPC1 | KD | HepG2 | ENCSR910YNJ | ENCSR279HMU |
| PABPC1 | KD | K562 | ENCSR192GBD | ENCSR419JMU |
| PABPC4 | KD | HepG2 | ENCSR455VZH | ENCSR237YZT |
| PABPC4 | KD | K562 | ENCSR047EEG | ENCSR344XID |
| PABPN1 | KD | K562 | ENCSR416ZJH | ENCSR164MUK |
| PAPOLA | KD | K562 | ENCSR368ZRP | ENCSR942UNX |
| PARN | KD | HepG2 | ENCSR825QXH | ENCSR305XWT |
| PARN | KD | K562 | ENCSR306IOF | ENCSR667PLJ |
| PARP1 | KO | HepG2 | ENCSR202SRN | ENCSR750CPT |
| PARP1 | KO | K562 | ENCSR226HQD | ENCSR712CQC |
| PCBP1 | KD | HepG2 | ENCSR635FRH | ENCSR603TCV |
| PCBP1 | KD | K562 | ENCSR545AIK | ENCSR129RWD |
| PCBP2 | KD | HepG2 | ENCSR028ITN | ENCSR491FOC |
| PCBP2 | KD | K562 | ENCSR648QFY | ENCSR129RWD |
| PCBP3 | KO | K562 | ENCSR105LOV | ENCSR341TTW |
| PDCD4 | KO | HepG2 | ENCSR580XTY | ENCSR670KAS |
| PDCD4 | KO | K562 | ENCSR469UOO | ENCSR516TLX |
| PES1 | KD | HepG2 | ENCSR496ETJ | ENCSR538QOG |
| PES1 | KD | K562 | ENCSR891DYO | ENCSR661HEL |
| PES1 | KO | K562 | ENCSR717XCO | ENCSR292PXV |
| PHF6 | KD | HepG2 | ENCSR681SMT | ENCSR674KEK |

**Table S3 – continued**

| | | | | |
|---|---|---|---|---|
| PHF6 | KD | K562 | ENCSR322XVS | ENCSR572FFX |
| PKM | KD | HepG2 | ENCSR656DQV | ENCSR689PHN |
| PKM | KD | K562 | ENCSR978CSQ | ENCSR438MDN |
| PNPT1 | KD | HepG2 | ENCSR880DEH | ENCSR279HMU |
| PNPT1 | KD | K562 | ENCSR191VWK | ENCSR620PUP |
| POLR2G | KD | K562 | ENCSR936TED | ENCSR942UNX |
| PPIG | KD | HepG2 | ENCSR620HAA | ENCSR246RRQ |
| PPIG | KD | K562 | ENCSR529MBZ | ENCSR032YMP |
| PPIL4 | KD | HepG2 | ENCSR851KEX | ENCSR305XWT |
| PPIL4 | KD | K562 | ENCSR556FNN | ENCSR419JMU |
| PPP1R10 | KO | HepG2 | ENCSR695RQF | ENCSR670KAS |
| PPP1R10 | KO | K562 | ENCSR439CUH | ENCSR516TLX |
| PPP1R8 | KD | K562 | ENCSR844QNT | ENCSR164MUK |
| PPP1R8 | KO | HepG2 | ENCSR592AQT | ENCSR827PII |
| PRPF3 | KO | HepG2 | ENCSR909TFJ | ENCSR704VPL |
| PRPF3 | KO | K562 | ENCSR681ELF | ENCSR877JEM |
| PRPF39 | KO | HepG2 | ENCSR511QFJ | ENCSR670KAS |
| PRPF4 | KD | K562 | ENCSR363QIA | ENCSR260BQC |
| PRPF6 | KD | HepG2 | ENCSR529QEZ | ENCSR491FOC |
| PRPF6 | KD | K562 | ENCSR783LUA | ENCSR815CVQ |
| PRPF8 | KD | HepG2 | ENCSR998MZP | ENCSR856ZRV |
| PRPF8 | KD | K562 | ENCSR137HKS | ENCSR661HEL |
| PRPF8 | KO | K562 | ENCSR320IEK | ENCSR015KGT |
| PSIP1 | KD | HepG2 | ENCSR744YVR | ENCSR674KEK |
| PSIP1 | KD | K562 | ENCSR611LQB | ENCSR572FFX |
| PSMA1 | KO | HepG2 | ENCSR608CUI | ENCSR953VMB |
| PSMA1 | KO | K562 | ENCSR058MCC | ENCSR712CQC |
| PTBP1 | KD | HepG2 | ENCSR064DXG | ENCSR603TCV |

**Table S3 – continued**

| | | | | |
|---|---|---|---|---|
| PTBP1 | KD | K562 | ENCSR527IVX | ENCSR129RWD |
| PTGES3 | KO | HepG2 | ENCSR528LDT | ENCSR964HKT |
| PTGES3 | KO | K562 | ENCSR728VRY | ENCSR404OHQ |
| PUF60 | KD | HepG2 | ENCSR648BSC | ENCSR042QTH |
| PUF60 | KD | K562 | ENCSR558XNA | ENCSR620PUP |
| PUM1 | KD | HepG2 | ENCSR945XKW | ENCSR042QTH |
| PUM1 | KD | K562 | ENCSR745WVZ | ENCSR620PUP |
| PUM2 | KD | HepG2 | ENCSR210DML | ENCSR042QTH |
| PUM2 | KD | K562 | ENCSR118XYK | ENCSR620PUP |
| PUS1 | KD | HepG2 | ENCSR296ERI | ENCSR538QOG |
| PUS1 | KD | K562 | ENCSR618IQH | ENCSR143COQ |
| QKI | KD | HepG2 | ENCSR330YOU | ENCSR603TCV |
| QKI | KD | K562 | ENCSR256PLH | ENCSR344XID |
| RAE1 | KO | HepG2 | ENCSR723QPD | ENCSR670KAS |
| RAE1 | KO | K562 | ENCSR092QTF | ENCSR516TLX |
| RAVER1 | KD | HepG2 | ENCSR576GOW | ENCSR491FOC |
| RAVER1 | KD | K562 | ENCSR904BCZ | ENCSR344XID |
| RBFOX2 | KD | HepG2 | ENCSR767LLP | ENCSR104ABF |
| RBFOX2 | KD | K562 | ENCSR336DFS | ENCSR667PLJ |
| RBM14 | KO | HepG2 | ENCSR166MWM | ENCSR521WAI |
| RBM14 | KO | K562 | ENCSR821KIN | ENCSR347NKX |
| RBM15 | KD | HepG2 | ENCSR599PXD | ENCSR585KOJ |
| RBM15 | KD | K562 | ENCSR385UPQ | ENCSR143COQ |
| RBM17 | KD | HepG2 | ENCSR385TMY | ENCSR104ABF |
| RBM17 | KD | K562 | ENCSR898OPN | ENCSR118EFE |
| RBM19 | KO | HepG2 | ENCSR532IHF | ENCSR306EWE |
| RBM19 | KO | K562 | ENCSR129AAZ | ENCSR978MMZ |
| RBM22 | KD | HepG2 | ENCSR330KHN | ENCSR997HCQ |

**Table S3 – continued**

| | | | | |
|---|---|---|---|---|
| RBM22 | KD | K562 | ENCSR947OIM | ENCSR129RWD |
| RBM25 | KD | HepG2 | ENCSR610AEI | ENCSR603TCV |
| RBM25 | KD | K562 | ENCSR149DMY | ENCSR344XID |
| RBM25 | KO | HepG2 | ENCSR739ZMF | ENCSR232ERY |
| RBM25 | KO | K562 | ENCSR041IXV | ENCSR347NKX |
| RBM26 | KO | K562 | ENCSR096JMV | ENCSR347NKX |
| RBM27 | KD | HepG2 | ENCSR222SMI | ENCSR674KEK |
| RBM28 | KO | K562 | ENCSR519LZD | ENCSR347NKX |
| RBM3 | KD | K562 | ENCSR675KPR | ENCSR936VPP |
| RBM34 | KD | HepG2 | ENCSR318HAT | ENCSR491FOC |
| RBM34 | KD | K562 | ENCSR560AYQ | ENCSR129RWD |
| RBM39 | KD | HepG2 | ENCSR760EGM | ENCSR104ABF |
| RBM39 | KD | K562 | ENCSR678WOA | ENCSR164MUK |
| RBM47 | KD | HepG2 | ENCSR711ZJQ | ENCSR279HMU |
| RBM5 | KO | HepG2 | ENCSR606PVX | ENCSR481WJH |
| RBM8A | KO | K562 | ENCSR163RXI | ENCSR347NKX |
| RCC2 | KD | HepG2 | ENCSR685JXU | ENCSR674KEK |
| RCC2 | KD | K562 | ENCSR921KDS | ENCSR572FFX |
| RECQL | KD | HepG2 | ENCSR572AMC | ENCSR603TCV |
| RECQL | KD | K562 | ENCSR310VND | ENCSR344XID |
| REXO2 | KO | HepG2 | ENCSR199WMC | ENCSR232ERY |
| RIOK1 | KO | K562 | ENCSR608WYZ | ENCSR347NKX |
| RNASEH2A | KO | HepG2 | ENCSR703FPC | ENCSR232ERY |
| RNASEH2A | KO | K562 | ENCSR002JOW | ENCSR347NKX |
| RNF187 | KO | HepG2 | ENCSR637CZY | ENCSR306EWE |
| RNF187 | KO | K562 | ENCSR050ONB | ENCSR978MMZ |
| RNH1 | KO | HepG2 | ENCSR706QGC | ENCSR232ERY |
| RNPC3 | KO | K562 | ENCSR631NIZ | ENCSR347NKX |

**Table S3 – continued**

| | | | | |
|---|---|---|---|---|
| ROCK2 | KO | HepG2 | ENCSR034VEC | ENCSR232ERY |
| RPL11 | KO | HepG2 | ENCSR702LIH | ENCSR964HKT |
| RPL11 | KO | K562 | ENCSR816LHY | ENCSR404OHQ |
| RPL18 | KO | HepG2 | ENCSR468LEL | ENCSR704VPL |
| RPL23A | KD | HepG2 | ENCSR706SXN | ENCSR674KEK |
| RPL23A | KO | HepG2 | ENCSR925OFV | ENCSR300EBW |
| RPL23A | KO | K562 | ENCSR997ACY | ENCSR015KGT |
| RPL28 | KO | HepG2 | ENCSR687ITW | ENCSR670KAS |
| RPL28 | KO | K562 | ENCSR234FWR | ENCSR404OHQ |
| RPL29 | KO | HepG2 | ENCSR333JNM | ENCSR232ERY |
| RPL29 | KO | K562 | ENCSR036ZTN | ENCSR347NKX |
| RPLP0 | KD | HepG2 | ENCSR082YGI | ENCSR674KEK |
| RPLP0 | KO | HepG2 | ENCSR000SKS | ENCSR105NML |
| RPLP0 | KO | K562 | ENCSR928MEW | ENCSR292PXV |
| RPRD1B | KO | HepG2 | ENCSR747MHD | ENCSR232ERY |
| RPRD1B | KO | K562 | ENCSR737OPD | ENCSR347NKX |
| RPS10 | KD | HepG2 | ENCSR410UHJ | ENCSR491FOC |
| RPS10 | KD | K562 | ENCSR004RGI | ENCSR084SCN |
| RPS11 | KO | K562 | ENCSR050ATA | ENCSR015KGT |
| RPS15A | KO | HepG2 | ENCSR119RXN | ENCSR232ERY |
| RPS15A | KO | K562 | ENCSR292NHK | ENCSR347NKX |
| RPS19 | KD | HepG2 | ENCSR486AIO | ENCSR689PHN |
| RPS19 | KD | K562 | ENCSR098NHI | ENCSR661HEL |
| RPS2 | KD | HepG2 | ENCSR667RIA | ENCSR674KEK |
| RPS21 | KO | K562 | ENCSR925SFP | ENCSR271ZAB |
| RPS25 | KO | HepG2 | ENCSR565ZPZ | ENCSR306EWE |
| RPS28 | KO | K562 | ENCSR928XFZ | ENCSR271ZAB |
| RPS3 | KD | K562 | ENCSR410MIQ | ENCSR084SCN |

**Table S3 – continued**

| | | | | |
|---|---|---|---|---|
| RPS3A | KD | HepG2 | ENCSR118VQR | ENCSR279HMU |
| RPS3A | KD | K562 | ENCSR788YGG | ENCSR419JMU |
| RPS5 | KD | HepG2 | ENCSR838SMC | ENCSR674KEK |
| RPS6 | KO | K562 | ENCSR539PFV | ENCSR712CQC |
| RPS8 | KO | HepG2 | ENCSR994NGU | ENCSR452TFV |
| RPUSD2 | KO | HepG2 | ENCSR235JSA | ENCSR452TFV |
| RPUSD2 | KO | K562 | ENCSR563QCQ | ENCSR271ZAB |
| RRP9 | KD | HepG2 | ENCSR471GIS | ENCSR856ZRV |
| RRP9 | KD | K562 | ENCSR210KJB | ENCSR661HEL |
| RTCB | KO | HepG2 | ENCSR726RTM | ENCSR452TFV |
| RTF1 | KD | HepG2 | ENCSR906WTM | ENCSR491FOC |
| RTF1 | KD | K562 | ENCSR783YSQ | ENCSR815CVQ |
| RYBP | KO | HepG2 | ENCSR439TMT | ENCSR306EWE |
| RYBP | KO | K562 | ENCSR607RGV | ENCSR978MMZ |
| SAFB | KO | HepG2 | ENCSR416YFB | ENCSR300EBW |
| SAFB | KO | K562 | ENCSR336TYW | ENCSR341TTW |
| SAFB2 | KD | HepG2 | ENCSR110ZYD | ENCSR305XWT |
| SAFB2 | KD | K562 | ENCSR770LYW | ENCSR154OBA |
| SARNP | KO | HepG2 | ENCSR207GTU | ENCSR452TFV |
| SARNP | KO | K562 | ENCSR639XFV | ENCSR271ZAB |
| SART1 | KO | HepG2 | ENCSR400YKV | ENCSR704VPL |
| SART1 | KO | K562 | ENCSR208AJS | ENCSR854OFV |
| SART3 | KD | HepG2 | ENCSR011BBS | ENCSR279HMU |
| SART3 | KD | K562 | ENCSR954HAY | ENCSR419JMU |
| SBDS | KD | HepG2 | ENCSR343DHN | ENCSR279HMU |
| SBDS | KD | K562 | ENCSR219DXZ | ENCSR419JMU |
| SCAF8 | KO | HepG2 | ENCSR330VES | ENCSR452TFV |
| SCAF8 | KO | K562 | ENCSR329UEC | ENCSR271ZAB |

**Table S3 – continued**

| | | | | |
|---|---|---|---|---|
| SEC23IP | KO | HepG2 | ENCSR745KNE | ENCSR452TFV |
| SEC63 | KO | HepG2 | ENCSR981TQN | ENCSR452TFV |
| SEC63 | KO | K562 | ENCSR082BCP | ENCSR271ZAB |
| SERBP1 | KD | HepG2 | ENCSR820ROH | ENCSR689PHN |
| SERBP1 | KD | K562 | ENCSR925RNE | ENCSR661HEL |
| SF1 | KD | HepG2 | ENCSR644AIM | ENCSR104ABF |
| SF1 | KD | K562 | ENCSR562CCA | ENCSR164MUK |
| SF3A1 | KO | HepG2 | ENCSR556TJU | ENCSR670KAS |
| SF3A1 | KO | K562 | ENCSR547ICR | ENCSR516TLX |
| SF3A2 | KO | HepG2 | ENCSR402BZZ | ENCSR671FDZ |
| SF3A2 | KO | K562 | ENCSR095TFW | ENCSR854OFV |
| SF3A3 | KD | HepG2 | ENCSR374NMJ | ENCSR691IVR |
| SF3A3 | KD | K562 | ENCSR454KYR | ENCSR174OYC |
| SF3B1 | KD | HepG2 | ENCSR896CFV | ENCSR264TUE |
| SF3B1 | KD | K562 | ENCSR047QHX | ENCSR424QCW |
| SF3B1 | KO | K562 | ENCSR544ZCR | ENCSR292PXV |
| SF3B2 | KO | K562 | ENCSR447ARO | ENCSR271ZAB |
| SF3B4 | KD | HepG2 | ENCSR148MQK | ENCSR603TCV |
| SF3B4 | KD | K562 | ENCSR081XRA | ENCSR344XID |
| SFPQ | KD | HepG2 | ENCSR782MXN | ENCSR104ABF |
| SFPQ | KD | K562 | ENCSR535YPK | ENCSR164MUK |
| SKIV2L2 | KO | HepG2 | ENCSR780UNO | ENCSR452TFV |
| SKIV2L2 | KO | K562 | ENCSR300UBK | ENCSR271ZAB |
| SLBP | KD | HepG2 | ENCSR519KXM | ENCSR042QTH |
| SLBP | KD | K562 | ENCSR112YTD | ENCSR667PLJ |
| SLTM | KD | HepG2 | ENCSR185JGT | ENCSR491FOC |
| SLTM | KD | K562 | ENCSR234YMW | ENCSR084SCN |
| SMN1 | KD | HepG2 | ENCSR090UMI | ENCSR305XWT |

**Table S3 – continued**

| | | | | |
|---|---|---|---|---|
| SMN1 | KD | K562 | ENCSR129ROE | ENCSR419JMU |
| SMNDC1 | KD | HepG2 | ENCSR995ZGJ | ENCSR491FOC |
| SMNDC1 | KD | K562 | ENCSR408SDL | ENCSR815CVQ |
| SMYD3 | KO | HepG2 | ENCSR989GNV | ENCSR306EWE |
| SMYD3 | KO | K562 | ENCSR775USN | ENCSR978MMZ |
| SND1 | KD | HepG2 | ENCSR398LZW | ENCSR491FOC |
| SND1 | KD | K562 | ENCSR232XRZ | ENCSR084SCN |
| SNRNP200 | KD | HepG2 | ENCSR003LSA | ENCSR104ABF |
| SNRNP200 | KD | K562 | ENCSR943LIB | ENCSR164MUK |
| SNRNP70 | KD | HepG2 | ENCSR635BOO | ENCSR264TUE |
| SNRPC | KO | HepG2 | ENCSR005RUR | ENCSR704VPL |
| SNRPC | KO | K562 | ENCSR840JRU | ENCSR877JEM |
| SPATS2 | KO | K562 | ENCSR535TPQ | ENCSR271ZAB |
| SRA1 | KO | K562 | ENCSR033MIV | ENCSR271ZAB |
| SRFBP1 | KD | HepG2 | ENCSR153GKS | ENCSR997HCQ |
| SRFBP1 | KD | K562 | ENCSR813NZP | ENCSR572FFX |
| SRP68 | KD | HepG2 | ENCSR167JPY | ENCSR856ZRV |
| SRP68 | KD | K562 | ENCSR312SRB | ENCSR661HEL |
| SRPK2 | KD | K562 | ENCSR524YXQ | ENCSR174OYC |
| SRSF1 | KD | HepG2 | ENCSR094KBY | ENCSR603TCV |
| SRSF1 | KD | K562 | ENCSR066VOO | ENCSR129RWD |
| SRSF3 | KD | HepG2 | ENCSR376FGR | ENCSR264TUE |
| SRSF3 | KO | K562 | ENCSR914IEV | ENCSR015KGT |
| SRSF4 | KD | K562 | ENCSR697GLD | ENCSR815CVQ |
| SRSF4 | KO | HepG2 | ENCSR471INA | ENCSR853AOV |
| SRSF4 | KO | K562 | ENCSR939BTN | ENCSR005JBU |
| SRSF5 | KD | HepG2 | ENCSR781YNI | ENCSR042QTH |
| SRSF5 | KD | K562 | ENCSR906RHU | ENCSR129RWD |

**Table S3 – continued**

| | | | | |
|---|---|---|---|---|
| SRSF5 | KO | K562 | ENCSR528YVC | ENCSR292PXV |
| SRSF7 | KD | HepG2 | ENCSR017PRS | ENCSR603TCV |
| SRSF7 | KD | K562 | ENCSR464ADT | ENCSR129RWD |
| SRSF7 | KO | HepG2 | ENCSR845MNM | ENCSR105NML |
| SRSF7 | KO | K562 | ENCSR139LOF | ENCSR163JUC |
| SRSF9 | KD | HepG2 | ENCSR597XHH | ENCSR603TCV |
| SRSF9 | KD | K562 | ENCSR113HRG | ENCSR129RWD |
| SRSF9 | KO | HepG2 | ENCSR814JQP | ENCSR300EBW |
| SRSF9 | KO | K562 | ENCSR972AZD | ENCSR341TTW |
| SSB | KD | HepG2 | ENCSR278CHI | ENCSR997HCQ |
| SSB | KD | K562 | ENCSR891AXF | ENCSR084SCN |
| SSRP1 | KD | HepG2 | ENCSR422JMS | ENCSR246RRQ |
| SSRP1 | KD | K562 | ENCSR902WSK | ENCSR032YMP |
| STAU1 | KD | HepG2 | ENCSR124KCF | ENCSR067GHD |
| STAU1 | KD | K562 | ENCSR777EDL | ENCSR032YMP |
| STAU2 | KO | HepG2 | ENCSR279MNU | ENCSR775ZWO |
| STAU2 | KO | K562 | ENCSR898KNS | ENCSR292PXV |
| STIP1 | KD | HepG2 | ENCSR871BXO | ENCSR279HMU |
| STIP1 | KD | K562 | ENCSR082UWF | ENCSR419JMU |
| SUB1 | KD | HepG2 | ENCSR997FOT | ENCSR491FOC |
| SUB1 | KD | K562 | ENCSR047AJA | ENCSR815CVQ |
| SUCLG1 | KD | HepG2 | ENCSR810JYX | ENCSR042QTH |
| SUCLG1 | KD | K562 | ENCSR101OPF | ENCSR620PUP |
| SUGP2 | KD | HepG2 | ENCSR837QDN | ENCSR246RRQ |
| SUGP2 | KD | K562 | ENCSR192BPV | ENCSR164MUK |
| SUPT6H | KD | HepG2 | ENCSR281KCL | ENCSR067GHD |
| SUPT6H | KD | K562 | ENCSR530BOP | ENCSR164MUK |
| SUPV3L1 | KD | HepG2 | ENCSR995RPB | ENCSR856ZRV |

**Table S3 – continued**

| | | | | |
|---|---|---|---|---|
| SUPV3L1 | KD | K562 | ENCSR778SIU | ENCSR661HEL |
| SYNCRIP | KO | K562 | ENCSR680OJB | ENCSR341TTW |
| TAF15 | KD | HepG2 | ENCSR998RZI | ENCSR603TCV |
| TAF15 | KD | K562 | ENCSR611ZAL | ENCSR344XID |
| TARDBP | KD | HepG2 | ENCSR527QNC | ENCSR264TUE |
| TARDBP | KD | K562 | ENCSR134JRE | ENCSR129RWD |
| TBRG4 | KD | HepG2 | ENCSR741YCA | ENCSR246RRQ |
| TBRG4 | KD | K562 | ENCSR079IPT | ENCSR344XID |
| TFIP11 | KD | HepG2 | ENCSR573UBF | ENCSR856ZRV |
| TFIP11 | KD | K562 | ENCSR911DGK | ENCSR661HEL |
| THOC1 | KO | HepG2 | ENCSR636PNL | ENCSR964HKT |
| THOC1 | KO | K562 | ENCSR921RQX | ENCSR404OHQ |
| TIA1 | KD | HepG2 | ENCSR057GCF | ENCSR603TCV |
| TIA1 | KD | K562 | ENCSR694LKY | ENCSR129RWD |
| TIAL1 | KD | HepG2 | ENCSR450VQO | ENCSR305XWT |
| TIAL1 | KD | K562 | ENCSR927TSP | ENCSR572FFX |
| TRA2A | KD | HepG2 | ENCSR030GZQ | ENCSR491FOC |
| TRA2A | KD | K562 | ENCSR916WOI | ENCSR129RWD |
| TRA2A | KO | K562 | ENCSR564RKL | ENCSR292PXV |
| TRIM56 | KD | HepG2 | ENCSR309HXK | ENCSR305XWT |
| TRIM56 | KD | K562 | ENCSR300QFQ | ENCSR572FFX |
| TRIP6 | KD | K562 | ENCSR152IWT | ENCSR032YMP |
| TRNAU1AP | KO | HepG2 | ENCSR481AIB | ENCSR775ZWO |
| TRNAU1AP | KO | K562 | ENCSR889MBF | ENCSR341TTW |
| TROVE2 | KD | HepG2 | ENCSR946OFN | ENCSR689PHN |
| TROVE2 | KD | K562 | ENCSR060KRD | ENCSR092WKG |
| TUFM | KD | HepG2 | ENCSR459EMR | ENCSR042QTH |
| TUFM | KD | K562 | ENCSR602AWR | ENCSR620PUP |

**Table S3 – continued**

| | | | | |
|---|---|---|---|---|
| U2AF1 | KD | HepG2 | ENCSR372UWV | ENCSR067GHD |
| U2AF1 | KD | K562 | ENCSR342EDG | ENCSR344XID |
| U2AF2 | KD | HepG2 | ENCSR426UUG | ENCSR104ABF |
| U2AF2 | KD | K562 | ENCSR904CJQ | ENCSR344XID |
| UBE2L3 | KD | HepG2 | ENCSR424JSU | ENCSR042QTH |
| UBE2L3 | KD | K562 | ENCSR362XMY | ENCSR620PUP |
| UCHL5 | KD | HepG2 | ENCSR684HTV | ENCSR856ZRV |
| UCHL5 | KD | K562 | ENCSR678MVE | ENCSR661HEL |
| UPF1 | KD | HepG2 | ENCSR689MIY | ENCSR691IVR |
| UPF1 | KD | K562 | ENCSR251ABP | ENCSR174OYC |
| UPF2 | KD | HepG2 | ENCSR318OXM | ENCSR856ZRV |
| UPF2 | KD | K562 | ENCSR810FHY | ENCSR438MDN |
| UTP18 | KD | HepG2 | ENCSR269ZAO | ENCSR691IVR |
| UTP18 | KD | K562 | ENCSR165VBD | ENCSR143COQ |
| UTP3 | KD | HepG2 | ENCSR910ECL | ENCSR691IVR |
| WBP11 | KO | HepG2 | ENCSR484QEX | ENCSR704VPL |
| WBP11 | KO | K562 | ENCSR070AYA | ENCSR877JEM |
| WDR3 | KD | HepG2 | ENCSR902VPO | ENCSR225PRV |
| WDR3 | KD | K562 | ENCSR334BTA | ENCSR174OYC |
| WDR43 | KD | K562 | ENCSR341PZW | ENCSR174OYC |
| WDR43 | KO | HepG2 | ENCSR606QVN | ENCSR827PII |
| WRN | KD | HepG2 | ENCSR093FHC | ENCSR225PRV |
| WRN | KD | K562 | ENCSR165BCF | ENCSR718EWL |
| XPO1 | KD | HepG2 | ENCSR438UOT | ENCSR225PRV |
| XPO1 | KD | K562 | ENCSR629XTS | ENCSR260BQC |
| XPO5 | KD | HepG2 | ENCSR778RWJ | ENCSR279HMU |
| XPO5 | KD | K562 | ENCSR453HKS | ENCSR164MUK |
| XRCC5 | KD | HepG2 | ENCSR732IYM | ENCSR491FOC |

**Table S3 – continued**

| | | | | |
|---|---|---|---|---|
| XRCC5 | KD | K562 | ENCSR715XZS | ENCSR815CVQ |
| XRCC6 | KD | HepG2 | ENCSR500WHE | ENCSR491FOC |
| XRCC6 | KD | K562 | ENCSR232CPD | ENCSR815CVQ |
| XRN1 | KD | HepG2 | ENCSR676SDU | ENCSR225PRV |
| XRN1 | KD | K562 | ENCSR949RMX | ENCSR260BQC |
| XRN2 | KD | HepG2 | ENCSR347ZHQ | ENCSR856ZRV |
| XRN2 | KD | K562 | ENCSR717SJA | ENCSR661HEL |
| YBX3 | KD | HepG2 | ENCSR494VSD | ENCSR305XWT |
| YBX3 | KD | K562 | ENCSR306EIU | ENCSR667PLJ |
| YTHDC2 | KD | HepG2 | ENCSR289HOV | ENCSR225PRV |
| YTHDC2 | KD | K562 | ENCSR843LYF | ENCSR164MUK |
| YWHAG | KD | HepG2 | ENCSR204QKA | ENCSR225PRV |
| YWHAG | KO | K562 | ENCSR163PNO | ENCSR015KGT |
| ZC3H11A | KO | HepG2 | ENCSR544LEZ | ENCSR775ZWO |
| ZC3H11A | KO | K562 | ENCSR048AUX | ENCSR292PXV |
| ZC3H8 | KD | K562 | ENCSR448JAM | ENCSR718EWL |
| ZNF106 | KO | HepG2 | ENCSR636DVA | ENCSR775ZWO |
| ZNF106 | KO | K562 | ENCSR660WRQ | ENCSR015KGT |
| ZNF622 | KD | HepG2 | ENCSR518JXY | ENCSR691IVR |
| ZRANB2 | KD | HepG2 | ENCSR081QQH | ENCSR491FOC |
| ZRANB2 | KD | K562 | ENCSR850PWM | ENCSR815CVQ |
